# A cytoskeletal scaffold promotes motile cilia assembly by regulating transition-zone integrity

**DOI:** 10.1101/2025.01.10.632296

**Authors:** Hyunji Park, Minjun Choi, Helen Oi-Lam Cheung, Yu Zhang, Shigeru Makino, Yoshiaki Yoshikawa, Haoran Qi, Zheng Liu, Guocheng Lan, Guoling Fu, Qian Wang, Shiny Shengzhen Guo, Pentao Liu, Zhen Liu, Shih-Chieh Ti, Won-Jing Wang, Xiang David Xiang Li, Tao Ni, Chi Chung Hui, Mu He

**Affiliations:** School of Biomedical Sciences, The University of Hong Kong, Hong Kong SAR, China; Program in Developmental & Stem Cell Biology, The Hospital for Sick Children, Toronto, ON M5G 0A4, Canada; Department of Molecular Genetics, University of Toronto, Toronto, ON M5S 1A8, Canada; Department of Chemistry, The University of Hong Kong, Hong Kong SAR, China; Department of Life Science, The Hong Kong University of Science and Technology, Hong Kong SAR, China; Center for Translational Stem Cell Biology, Hong Kong Science and Technology Park, Hong Kong SAR, China; Department of Molecular Medicine, Max Planck Institute of Biochemistry, Martinsried, Germany; Taiwan International Graduate Program in Molecular Medicine, National Yang Ming Chiao Tung University and Academia Sinica, Taipei, Taiwan; Institute of Biochemistry and Molecular Biology, National Yang Ming Chiao Tung University, Taipei 112, Taiwan

**Author notes:** These authors contribute equally to this work (Hyunji Park, Minjun Choi, Helen Oi-Lam Cheung).

## Abstract

Motile cilia are eukaryotic organelles with essential chemo- and mechano-sensing functions across evolution, from single cell organisms to humans. Motile cilia of the mammalian nervous, respiratory and reproductive systems are characterized by unique motility proteins to generate fluid flow essential for transporting metabolites and removing mucus. The molecular mechanism of motile cilia biogenesis remains unknown. Here, we use mouse genetics, single-molecule motility assays, proteomics, high-resolution imaging, and *in situ* cryo-tomography to identify mammalian KIF27, a motor protein of the Kinesin-4 family and homologue of the Hedgehog pathway regulator COS2/KIF7, as a key regulator of motile cilia assembly. We show that KIF27 promotes the integrity of the transition zone, a diffusion barrier situated at the cilium base. Loss of KIF27 causes specific and profound defects in axonemal structure and disrupts cilia beating, which collectively lead to organismal phenotypes that recapitulate primary ciliary dyskinesia. We show that the motile properties of KIF27 are dispensable for its function in motile cilia biogenesis. Instead, KIF27 acts as a microtubule scaffold to regulate the transition zone architecture and enable correct ciliary incorporation of motility-generating proteins. Given that KIF27 homologues exist in different evolutionarily lineages, we propose that the ancestral activities of KIF27/KIF7 kinesins were to form a microtubule-associated scaffold for protein-protein interactions pertinent to cilia formation and signaling. The transition-zone associated KIF27 activities may represent a general building principle for motile cilia assembly in diverse species and cell types.

## Introduction

Primary cilia and motile cilia are conserved microtubule-based organelles found in all modern eukaryotes and have diverse mechano-sensory functions. The primary cilium is well recognized as a hub of signaling pathways, including the hedgehog pathway^1,2^. Motile cilia, in contrast, can produce synchronized waveforms and drive extracellular fluid flow to regulate locomotion of marine larvae and transport mucus and metabolites across brain ventricles, respiratory and auditory epithelia, and reproductive tracks in mammals^3,4^. At the respiratory mucosal barrier, motile ciliated cells constitute the primary and foremost host defense mechanism by generating coordinated ciliary flow to remove mucus and attached pathogens^3,5^. Primary cilia and motile cilia dysfunctions can lead to ciliopathies and motile ciliopathies, respectively, with non-overlapping clinical presentations^3,6^. Primary ciliary dyskinesia (PCD) is a form of motile ciliopathy caused by genetic mutations that impair motile cilia beating. PCD patients exhibit chronic respiratory and ear inflammation, while some also experience situs inversus and subfertility^3,5,7^. Despite the importance of motile cilia to human physiology and disease, the molecular mechanisms that regulate motile cilia biogenesis are not well understood.

Both primary and motile cilia are compartmentalized from the cytosolic environment by unique membrane and cytoskeletal compositions to optimize for the specialized ciliary functions. Within cilia, axonemal microtubule doublets of a 9-fold symmetry are templated from modified centrioles, the basal bodies, which are docked onto the plasma membrane. During ciliogenesis, tubulin heterodimers, signaling molecules, and various ciliary structural components are transported by the intraflagellar trafficking (IFT) complex attached to the anterograde and retrograde motor proteins^8,9^. At the base of the cilia sits the transition zone (TZ), a sub-organellar domain comprised of conserved proteins, including CEP290, the Nephronophthisis (NPHP) complex, and the Meckel–Gruber syndrome (MKS) complex^10,11^. The organization of mammalian TZ in the primary cilium is well defined, and genetic studies demonstrate a gating mechanism for compartmentalization, in which the TZ regulates ciliary cargo entry and exit by acting as a critical diffusion barrier^12^. Many human ciliopathies are caused by mutations that perturb transition zone functions associated with the primary cilium. Motile cilia contain, in addition to the axonemal building blocks and TZ proteins, macromolecular machineries to drive coordinated cilia beating. These motility components include the central pair (CP) apparatus, radial spokes, outer and inner dynein arms, as well as a growing list of microtubule inner proteins (MIPs)^8,13,14^, which occupy the luminal space of axonemal microtubules and function to stabilize the axoneme and regulate motility^13,15,16^. Much of our understanding of motile cilia comes from studies in basal eukaryotes, such as *Chlamydomonas* and *Trypanosome brucei*. The molecular mechanisms that regulate motile cilia biogenesis and function in mammals have not been defined. It remains unknown how these motility proteins are incorporated into the ciliary compartment during motile cilia biogenesis, largely due to the lack of mammalian organismal and cell biological models for motile cilia formation and function.

To uncover the molecular mechanisms underlying the regulation of mammalian motile cilia assembly, we employed an *in-silico* strategy to identify potential components of the cilia biogenesis pathway. By combining mouse genetics, *in vitro* microtubule assays, cilia proteomics high-resolution cilia imaging, and *in situ* cryo-tomography, we found that an intact ciliary transition zone is essential for motile cilia assembly, and that formation of an intact transition zone depends on KIF27, a motor protein of the kinesin-4 family and homologue to the Hedgehog pathway regulator COS2/KIF7^17–21^. Loss of KIF27 causes specific and profound defects in axonemal structure and disrupts cilia beating, which collectively lead to organismal phenotypes that recapitulate PCD. Unexpectedly, KIF27 does not use its motor function for motile cilia biogenesis. Instead, KIF27 acts as a microtubule-associated scaffold to promote ciliary incorporation of motility proteins by controlling the structural integrity of the TZ. Given that KIF27 orthologues are present in many organisms with motile cilia, the transition-zone associated KIF27 activities may represent a general building principle for motile cilia assembly in diverse species and cell types.

## Results

### *In silico* enrichment for conserved motile cilia-associated genes

Motile cilia possess unique machineries to generate the “9+2” axonemal structure and motility (**Fig. 1a**). To identify novel regulators underlying motile cilia assembly, we carried out a stepwise *in-silico* curation to select for evolutionarily conserved genes specifically associated with motile ciliogenesis (**Fig. 1b**). Using publicly available single-cell RNA sequencing datasets, we first compiled a list of differentially expressed genes from motile ciliated cells of the mouse and human airways^22^, mouse ependyma^23^, as well as human fallopian tubes^24^. Next, we removed entries from ciliopathy and PCD gene panels^3,6^, as well as entries that encode proteins identified by proteomics from both primary and motile cilia^25^. Our filtering strategy led to about 80 genes (**Supplementary Table 1**), including *MAP9*^26^, *STK33*^27,28^, *KIF9*^29,30^, and *TTLL9*^31^, which are implicated in different aspects of motile cilia formation and function. From this list, we then performed text mining to search for entries with reported motile cilia phenotypes and prioritized those with orthologues in ciliated species ranging from *Trypanosoma*, sea anemone, to mice and humans (**Fig. 1b**). One of the entries that met all the criteria was *Kif27*, an uncharacterized motor protein that belongs to the Kinesin-4 family and is homologous to the Hh pathway regulators *Cos2/Kif7* (**Fig. 1c**). siRNA targeting Planaria *Kif27* disrupted ciliogenesis and locomotion^32^. A lacZ insertion to the mouse *Kif27* locus led to early lethality with hydrocephalus and nasal mucus accumulation^33^. These organismal defects in different species suggest an evolutionarily conserved role for *Kif27* in motile cilia.

**Figure 1.**
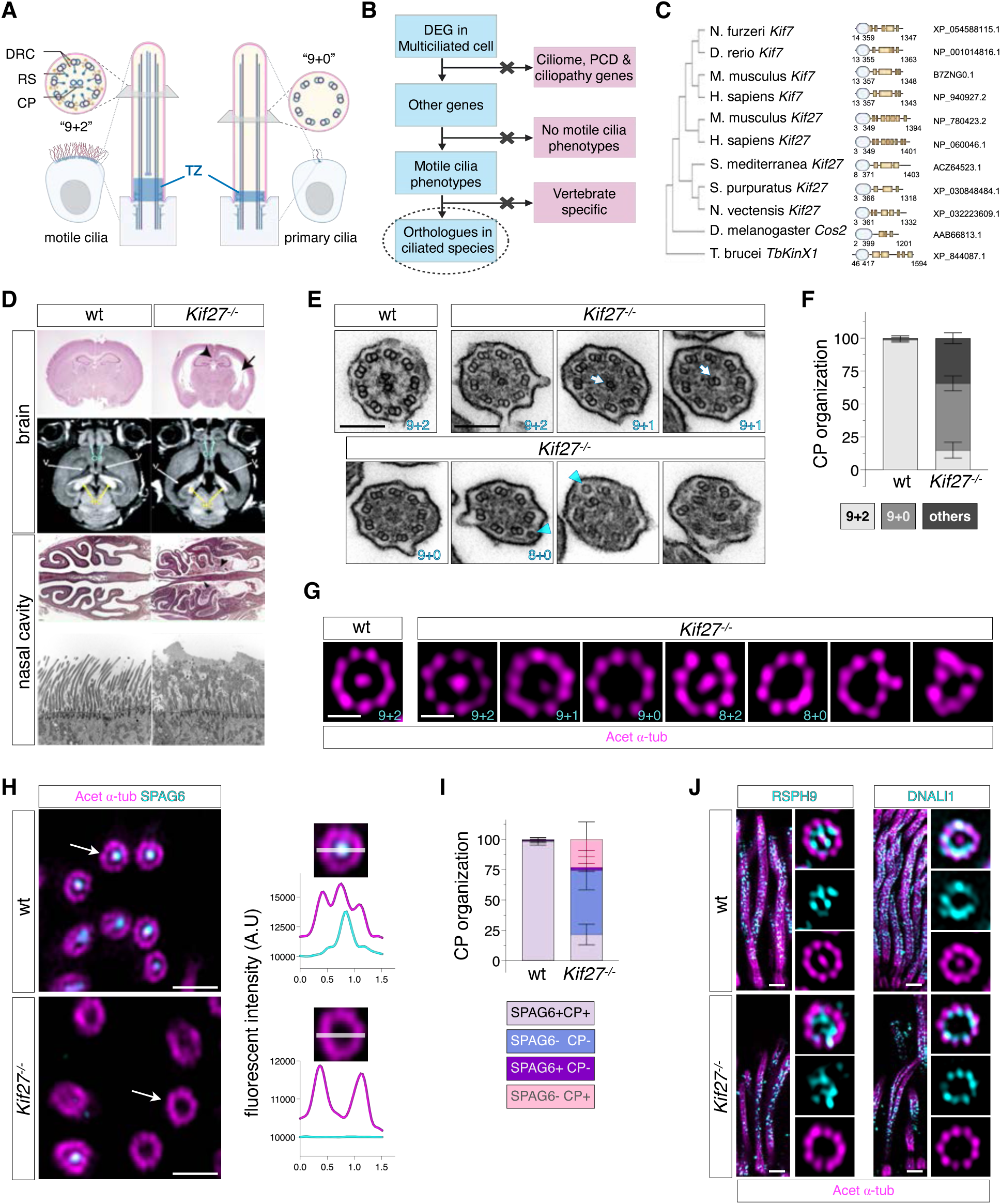
KIF27 is indispensable for mammalian motile ciliogenesis. (**A**) “9+2” motile cilia are comprised of 9 doublet microtubules surrounding a central pair (CP) microtubules and motility machineries including dynein regulatory complex (DRC) and radial spokes (RS). “9+0” primary cilia lack central pair and associated structures. The transition zone (TZ) sits at the cilium base. **(B)** Stepwise selection process for candidate gene identification. DEG: differentially expressed genes. **(C)** Left: Phylogenetic tree displaying homologs of *Kif27* in ciliated organisms. Middle: Domain architecture of *Kif27* homologs: kinesin motor domain (blue) and coiled-coil regions (yellow). Right: NCBI Reference sequences. **(D)** Histology and MRI analysis of mice brain reveal the enlarged ventricular zone (v) and compressed hippocampus (hc) and olfactory bulb (ob) were present in postnatal *Kif27^−/−^* mice compared to wild-type (wt). Loss of corpus callosum (arrowhead) and enlarged lateral ventricle (arrow) were also evident in mutants. Coronal section of *Kif27^−/−^*nasal olfactory epithelium show accumulation of mucus and dead cells in the nasal cavity (arrowheads). Ultrastructural analysis by transmission electron microscopy (TEM) in the nasal respiratory epithelium depicts asynchronized cilia array with trapped mucus between cilia. **(E)** TEM images of the respiratory cilia in postnatal day 17 (p17) wt and *Kif27^−/−^* mice. Cilia in wt show “9+2” axonemal configuration. Cilia in *Kif27^−/−^* exhibited axonemal organizations such as “9+1”, “9+0”, “8+0” and lacked two singlet CP microtubules (arrows). Others displayed loss of one or more peripheral doublets (arrowheads). Scale bars = 500 nm. **(F)** Quantification of ultrastructural defects in CP organization in wt trachea (n = 114, N = 4) and *Kif27^−/−^* trachea (n = 204, N = 4). Others include “9+1”, “9+0”, “8+0”, and disorganized axonemes. Error bars indicate standard deviation (SD). **(G)** Respiratory ciliary axonemes labeled with acetylated α-tubulin in magenta and observed with expansion microscopy p17 wt and *Kif27^−/−^* mice. Scale bars = 500 nm. **(H)** *En face* images of postnatal respiratory cilia in wt and *Kif27^−/−^* mice taken using tissue ultrastructure expansion microscopy. SPAG6 in cyan labels CP associated apparatus and acetylated α-tubulin in magenta marks axonemal microtubules. Line scan analyses for fluorescence intensities of SPAG6 and acetylated α-tubulin of indicated axonemes (arrows) for wt and mutants. Scale bar = 2 µm. **(I)** Quantification of CP and SPAG6 organization in p17 wt (n = 260, N = 3) and *Kif27^−/−^* mice (n = 260, N = 3). Error bars indicate SD. **(J)** The organization and expression of RSPH9 and DNALI1, both in cyan, in postnatal wt and *Kif27^−/−^* cilia. Acetylated α-tubulin in magenta marks axonemal microtubules. Scale bar = 1 µm.

### Kif27 is indispensable for motile cilia biogenesis, but not for Hh signaling

To investigate KIF27 function and molecular mechanism in mammalian motile cilia, we generated a *Kif27^−/−^* knockout mouse line by removing the first protein coding exon of *Kif27* (**Extended Data Fig. 1a-e**). *Kif27^−/−^* homozygous null mice exhibited hydrocephalus and nasal mucus accumulation (**Fig. 1d**), characteristics of PCD-related pathology. Using scanning electron microscopy (SEM), we observed reduced ciliation in the brain ependyma and morphological defects for respiratory motile cilia including reduced cilia length, supporting a role for KIF27 in motile cilia biogenesis (**Extended Data Fig. 2a-c**).

KIF27 is a paralogue of mammalian KIF7, which plays a dual role in Hh signaling by regulating primary cilia structure and controlling primary ciliary dynamics of the GLI transcription factor, the final effector of Hh signal transduction^34–36^. In contrast to *Kif7^−/−^* embryos which show gain-of-function Hh mediated patterning defects, *Kif27^−/−^* mice did not show detectable defects in Hh-dependent developmental processes (**Extended Data Fig. 3a-c**). *Kif27^−/−^*; *Kif7^−/−^* double mutants died at birth and exhibited polydactyly (**Extended Data Fig. 3b**), phenocopying the *Kif7^−/−^* single mutant defects. These genetic analyses provide evidence against a strong role for *Kif27* in primary cilia-mediated Hh signaling. Together, the data demonstrate a motile cilia-specific function for mammalian *KIF27*.

### Loss of KIF27 causes defects in motile cilia axonemal structure

Using transmission electron microscopy (TEM), we observed an array of axoneme abnormalities in postnatal *Kif27^−/−^* respiratory cilia. The normal “9+2” configuration of motile cilia with 9 outer doublets + 2 singlet central pair microtubules was only present in < 20% of *Kif27^−/−^* respiratory cilia (**Fig. 1e**). Instead, a fraction of axonemes showed either one singlet or doublet CP microtubules; nearly 40% of cilia completely lacked CP microtubules (9+0); about 30% of *Kif27^−/−^* cilia displayed < 9 outer doublets, open B-tubules, and disorganized doublets (**Fig. 1f**). Consistent with TEM analysis, tissue expansion microscopy with acetylated α-tubulin staining revealed striking cilia defects, including missing CP, reduction in outer doublets, as well as mis-arranged doublets in respiratory epithelia of postnatal *Kif27^−/−^* mice (**Fig. 1g**).

We next used expansion microscopy to characterize cilia defects in *Kif27^−/−^* mutants in more detail. SPAG6 is a CP apparatus protein associated with one of the CP singlet microtubules^37^. In both embryonic and postnatal *Kif27^−/−^* mice, > 70% reduction in SPAG6+ motile cilia were observed compared to wild-type (wt), suggesting that axonemal defects arise early in *Kif27^−/−^* mice (**Fig. 1h, i and Extended Data Fig. 2d-i**). In wt respiratory cilia, SPAG6 signal appeared as discrete puncta adjacent to the CP microtubule labeled with acetylated α-tubulin (**Fig. 1h and Extended Data Fig. 2h**). About 50% of *Kif27^−/−^* cilia lacked detectable signals for both SPAG6 and CP tubulin (SPAG6-CP−); about 25% of *Kif27^−/−^*cilia were SPAG6-CP+; 2% of *Kif27^−/−^* cilia were categorized as SPAG6+ CP−, suggesting that SPAG6 is unlikely to incorporate into the CP apparatus without a microtubule scaffold (**Fig. 1i and Extended Data Fig. 2i**). In contrast, RSPH9^38^ and DNALI1^39^, which are subunits of radial spokes and the inner dynein arm complex, respectively, were organized in the ciliary compartment similarly between wt and *Kif27^−/−^* cilia, regardless of the presence of CP (**Fig. 1j and Extended Data Fig. 2j**).

### Dynamic localization of KIF27 during motile cilia biogenesis

To characterize the endogenous KIF27 localization, we generated a KIF27-GFP reporter allele in which GFP was fused to the last exon of the mouse *Kif27* locus (**Extended Data Fig. 4a**). The homozygous *Kif27^GFP/GFP^* as well as *Kif27^GFP/−^*mice were viable and fertile, indicating that the KIF27-GFP fusion protein is functional and can inform the endogenous localization of KIF27 during different stages of motile ciliogenesis. In the embryonic respiratory ciliated cells of *Kif27^GFP/GFP^* animals, GFP signal was first detected in the deuterosome stage and co-localized with CP110^40,41^, a centrosomal protein, at the distal end of the amplifying centrioles prior to cilia formation (**Fig. 2a**). When centrioles were docked onto the apical plasma membrane, GFP signals co-localized with CEP164^42^, a cilia transition fiber marker (**Fig. 2b**). When ciliation was initiated, KIF27-GFP was enriched in the axonemal distal ends of newly formed short cilia but became undetectable in long and mature cilia (**Fig. 2c**). The dynamic associations of KIF27-GFP with centrioles and nascent ciliary axoneme suggest that KIF27 functions may be temporally regulated during ciliogenesis.

**Figure 2.**
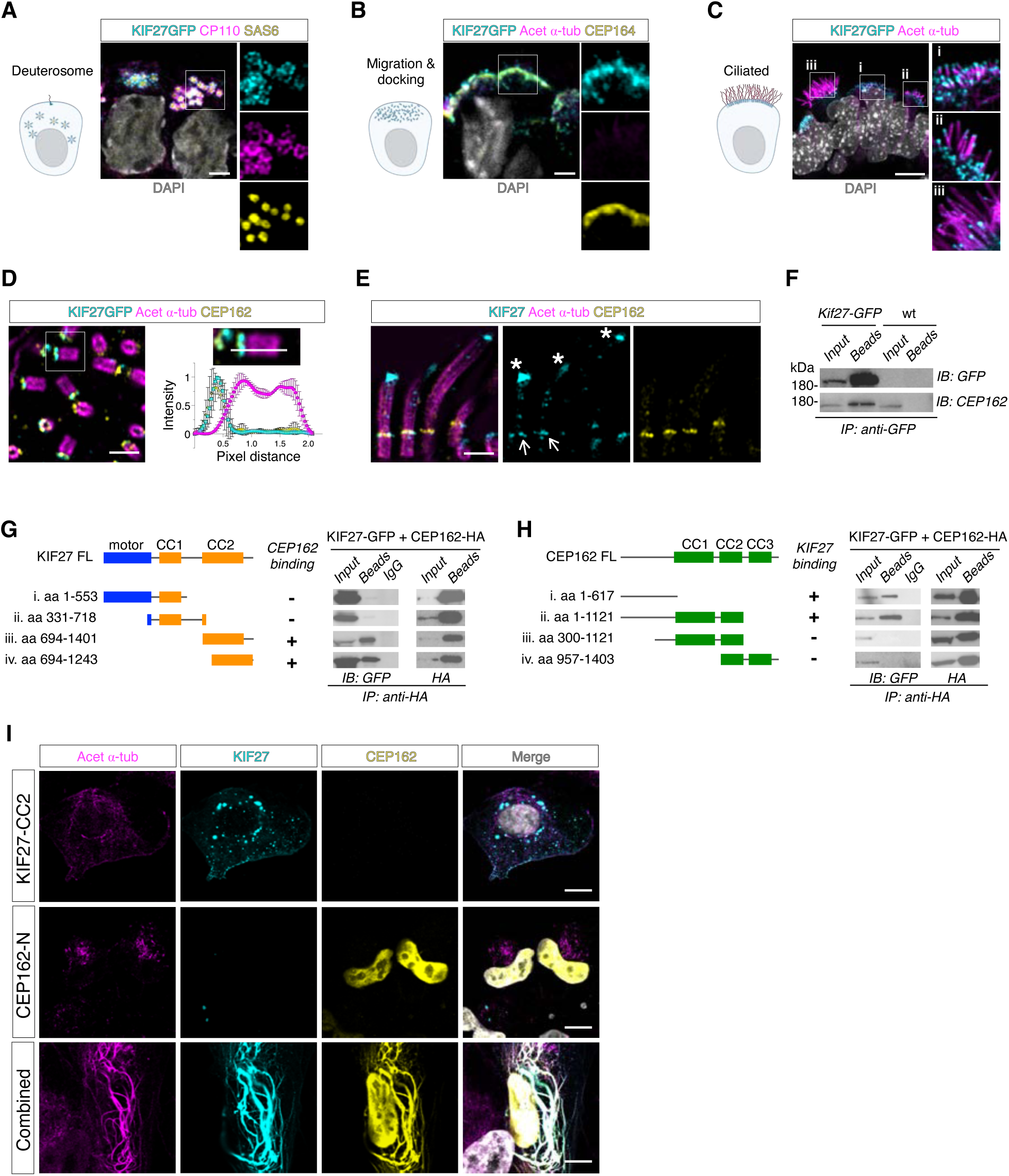
KIF27 localizes to motile cilia and interacts with CEP162. **(A)** Endogenous KIF27-GFP (cyan) co-localizes with distal centriole protein CP110 (magenta) during centriole amplification at the deuterosome stage of E16 *Kif27^GFP/GFP^* mouse nasal epithelium. SAS6 in yellow labels the centriole cartwheel. **(B)** KIF27-GFP (cyan) co-localizes with transition fiber protein CEP164 (yellow) when the centrioles are docked. Centriolar microtubules are labeled by acetylated α-tubulin in magenta; the signal was faint at this stage. **(C)** KIF27-GFP (cyan) localizes at the tips of the short cilia and becomes undetectable when cilia elongate. Cilia axonemes are labeled by labeled by acetylated α-tubulin in magenta. DAPI in grey labels the nucleus; scale bars = 2 µm for **(A)**-**(C)**. **(D)** KIF27-GFP (cyan) co-localizes with distal centriole protein CEP162 (yellow) during centriole amplification stage as shown by expansion microscopy and linescan analysis. Centriolar microtubules are labeled by acetylated α-tubulin in magenta. Scale bar = 500 nm. **(E)** KIF27 (cyan), labeled with a KIF27 antibody, co-localizes with CEP162 (yellow) at the transition zone (arrows) and at the tip of the short cilia (asterisks) in the embryonic wt respiratory epithelium. Scale bars = 2 µm. **(F)** Endogenous KIF27-GFP and CEP162 in *Kif27^GFP/GFP^* mouse testis co-immunoprecipitated in anti-GFP pull-down by GFP nanobody. Endogenous CEP162 remained in the unbound fraction when performing anti-GFP pull-down using wt (*Kif27 ^+/+^*) mouse testis. Input (cell lysate) and beads (eluate fraction) were subject to immunoblotting (IB) with anti-GFP and anti-CEP162 antibodies. **(G and H)** Left: Diagram of domain architecture of full-length human KIF27 **(G)** and full-length human CEP162 **(H)** and truncation constructs ((i)-(iv)). Right: anti-HA immunoprecipitation (IP) from HEK293T cells. Input (cell lysate), beads (eluate fraction) and IgG (eluate fraction from IgG isotype control) were subject to immunoblotting with anti-HA and anti-GFP antibodies. Positive and negative results of co-immunoprecipitation (Co-IP) of truncation construct with full-length protein are indicated as ‘+’ and ‘-’. **(I)** U2OS cells transiently expressing GFP-tagged KIF27-CC2 in cyan (aa 694-1401) or HA-tagged CEP162-N in yellow (aa 1-617) displaying cytoplasmic puncta or nuclear localization, respectively. Microtubule bundles present in U2OS cells transiently co-expressing GFP-tagged KIF27-CC2 and HA-tagged CEP162-N. Acetylated ⍺-tubulin in magenta labels microtubules. DAPI in grey labels the nucleus. Scale bar = 10 μm.

### KIF27 associates with transition zone protein CEP162 in motile ciliated cells

In human U2OS cells without motile cilia, CP110 was identified as a component of the centriole distal-end complex together with CEP97, CEP290, and CEP162^43^. CEP162, which binds to CEP290, is a microtubule-associated TZ protein^44^. Consistent with the co-localization of KIF27-GFP with CP110, we observed that in *Kif27^GFP/GFP^* mouse embryonic motile ciliated cells, KIF27-GFP co-localized with CEP162 at the distal end of amplifying centrioles (**Fig. 2d**). To validate the subcellular localizations of KIF27, we generated an antibody that binds to the central region of KIF27 (**Extended Data Fig. 1b-e**). Consistent with our KIF27-GFP analyses, antibody-labeled KIF27 specifically co-localized with CEP162 at the distal end of centrioles as well as the TZ in newly formed motile cilia, in which KIF27 was also enriched at the cilia tip (**Fig. 2e and Extended Data Fig. 5**).

To determine whether KIF27 can form a protein complex with CEP162, we purified a GFP nanobody to immunoprecipitate KIF27-GFP from mouse tissues and cells (**Extended Data Fig. 4b, c**). In tissue lysates from adult *Kif27^GFP/GFP^* testis, endogenous CEP162 co-immunoprecipitated with KIF27-GFP (**Fig. 2f**). To locate the interaction domains in each protein, structure-function analyses were performed by co-expressing truncation constructs of KIF27 and CEP162 (**Fig. 2g, h**). The binding domains were mapped to the C-terminus of KIF27 (KIF27-CC2) and the N-terminal flexible region of CEP162 (CEP162-N). The KIF27-binding domain of CEP162 is different from its microtubule-binding and CEP290-binding domains, which are located to the central coiled coils of CEP162^44^. To visualize the KIF27-CEP162 interaction in a cellular context, we expressed KIF27-GFP and HA-CEP162 binding constructs in U2OS cells. When expressed alone, KIF27-CC2 appeared in dispersed intracellular puncta, whereas CEP162-N showed a strong nuclear localization. In cells that co-expressed KIF27-CC2 and CEP162-N, both proteins showed translocation to cytoplasmic microtubule bundles (**Fig. 2i**). The results suggest that a physical association between KIF27 and CEP162 can induce microtubule binding of the protein complex, which may contribute to the stabilization of TZ microtubules and to confer mechanical rigidity at the cilia base.

### KIF27 is a motile kinesin with ATP dependent motility

Some members of the kinesin-4 family, for instance KIF4^45^ and KIF21^46,47^, show ATP dependent microtubule plus-end directed motility, while others, such as KIF7, lack ATP-mediated translocations along microtubules^36^. The ATP-dependent translocation ability of KIF27 has not been fully characterized. To address the molecular mechanism of KIF27 in motile cilia, we first characterized the biochemical properties of purified KIF27 motor proteins using well-established TIRF-microscopy assays^36,48^. A GFP tagged truncated KIF27 motor dimer, including the N-terminal motor domain and the first coiled-coil of KIF27, KIF27_1-570_-GFP, was purified from insect cells^35^ **(Fig. 3a and Extended Data Fig. 6a)**. To test the motor activity of KIF27 towards microtubules specific to motile cilia, we purified tubulin heterodimers TUBA1A-TUBB4B from insect cells^49,50^ and polymerized them into microtubules with GMP-CPP, a slowly hydrolyzing analog of GTP^48,51^ (**Extended Data Fig. 6b, c**). Translocations of GMPCPP-stabilized TUBA1A-TUBB4B microtubules were observed using TIRF assays in the presence of KIF27_1-570_GFP with 2mM Mg-ATP, but not with 0mM ATP or 2mM AMP-PNP, a non-hydrolysable analog of ATP (**Extended Data Fig. 7a**). We further characterized the motor properties of KIF27_1-570_GFP single particles using TIRF-microscopy assays (**Fig. 3b**). Analysis of kymographs revealed that in the presence of 1mM Mg-ATP, KIF27_1-570_GFP exhibited processive translocations along GMP-CPP stabilized TUBA1A-TUBB4B microtubules, with a mean velocity of 25.5 ± 1.2 nm/s and a mean microtubule binding of 38.8 ± 2.7 s (**Fig. 3c-e**). The *in vitro* velocity of KIF27 was orders of magnitude slower than KIF4^52^, indicating that KIF27 is unlikely to function as a ciliary transporter due to its slow-moving behavior.

**Figure 3.**
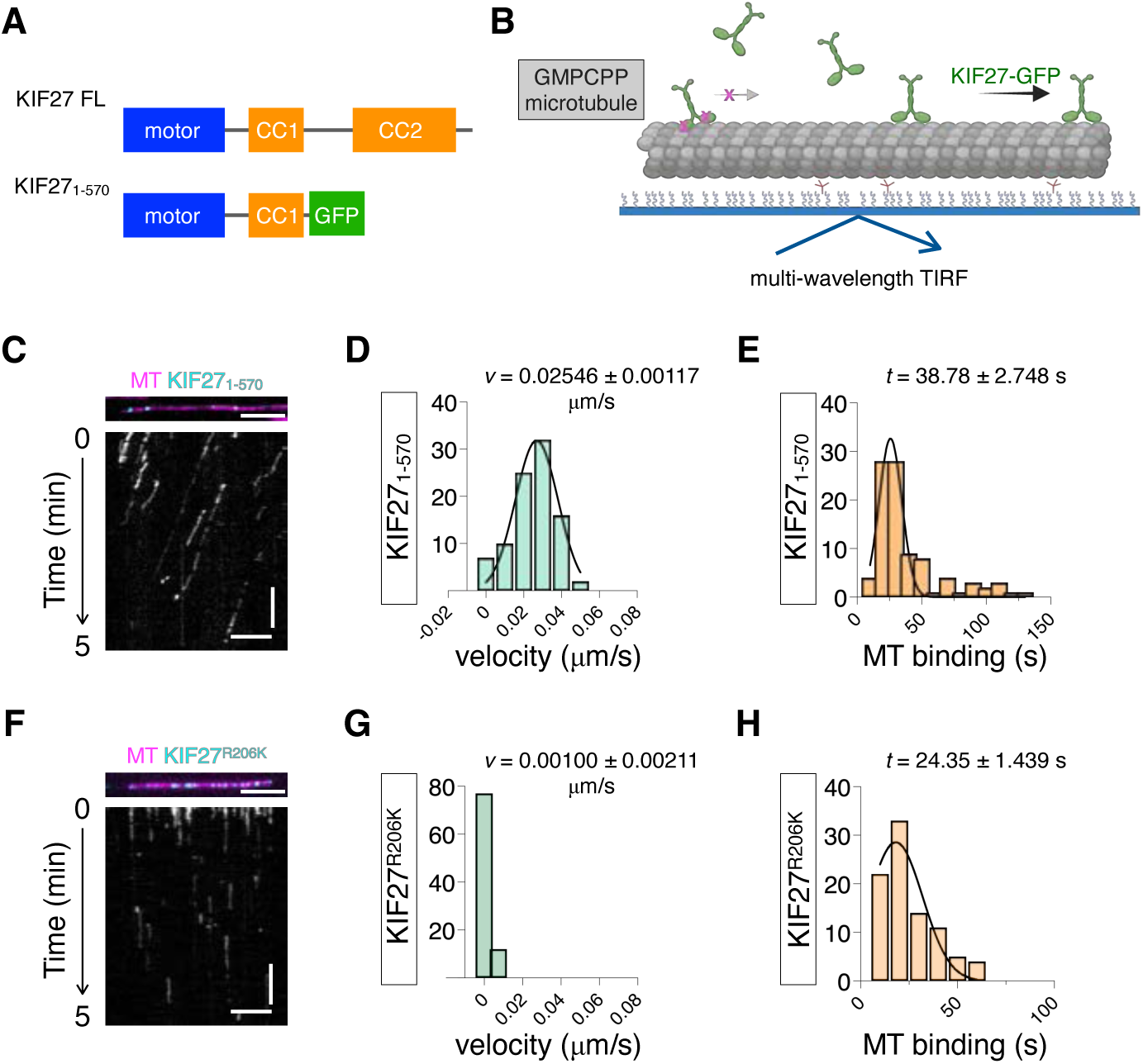
KIF27 is an ATP-dependent kinesin motor. (**A**) Domain organizations of full-length human KIF27_1-1401_, which includes the N-terminal kinesin motor domain and two coiled-coils, as well as the KIF27_1-570_-GFP motor dimer used in the in vitro motility assays. (**B**) Schematic representation of *in vitro* TIRF set-up for the motility assay. GMPCPP-stabilized microtubules (grey) were attached to a PEGylated coverslip (blue) by NeutrAvidin links before addition of active KIF27 motor proteins (green). The assay can determine the velocity and microtubule binding duration for KIF27-GFP motor constructs. **(C)** (top) Representative still image showing KIF27_1-570_-GFP (cyan) binding to X-rhodamine-labelled microtubules (magenta) in the presence of 2 mM ATP. Scale bar = 5 μm. (bottom) kymograph generated from the above KIF27 single molecule motility analysis. Horizontal scale bar = 5 µm. Vertical scale bar = 60 s. **(D and E)** Distribution of velocities **(D)** and binding durations **(E)** of KIF27_1-570_ (n = 92). (**F**) (top) Representative still image showing KIF27^R206K^-GFP (cyan) binding to X-rhodamine-labelled microtubules (magenta) in the presence of 2 mM ATP. Scale bar = 5 μm. (bottom) kymograph generated from the above KIF27^R206K^-GFP single molecule motility analysis. Horizontal scale bar = 5 µm. Vertical scale bar = 60 s. **(G and H)** Distribution of velocities **(G)** and binding durations **(H)** of KIF27^R206K^ (n = 89). The average velocity or MT binding duration is shown as mean ± standard error of mean.

### KIF27 ATP-dependent motility is dispensable for motile cilia biogenesis

*Drosophila* COS2 and mammalian KIF7 both act as microtubule associated scaffolds to mediate Hh signaling, raising the possibility that KIF27 may act as a scaffold in a motile cilia context. We next determined whether KIF27 ATP-dependent motility is required for its function in motile cilia. Two well-characterized mutations in the motor domain of the *Drosophila* kinesin heavy chain *Khc*, E164A^53–55^ and R210K^56,57^, can severely impair the ATP-dependent motility of the kinesin. E164 is located away from the ATP pocket of *Khc* and toward the motor-microtubule binding interface, and the E164A substitution results in a rigor kinesin that binds to ATP and microtubules with a higher affinity than wt. Khc R210 is within switch-I, a structural motif critical for ATP hydrolysis. Khc R210K homodimers lose the ability to hydrolyze ATP but remain associated with microtubules. Both E164 and R210 of *Drosophila Khc* are conserved in mammalian KIF27 (**Extended Data Fig. 7b, c**).

To test the effect of these substitutions on KIF27 ATP-dependent motor behaviors, we first expressed and purified the recombinant KIF27^E160A^ -GFP motor dimer analogous to the E164A *Drosophila Khc* mutant (**Extended Data Fig. 7d**). Using TIRF-microscopy assays, KIF27^E160A^ - GFP single particles showed a 36% reduction in motility and a 44% increase in microtubule binding when compared to the control KIF27_1-570_ motor dimers (**Extended Data Fig. 7f, g**). In contrast, the KIF27^R206K^-GFP, which contains the R206K substitution analogous to *Drosophila* R210K in *Khc* (**Extended Dat Fig. 7e**), completely blocked ATP-dependent KIF27 motility along TUBA1A-TUBB4B microtubules and resulted in a 40% decrease in microtubule binding duration (**Fig. 3f-h**).

Based on the *in vitro* biochemical properties of KIF27^R206K^ motor dimers, we generated a mouse *Kif27^R206K^* allele to investigate whether KIF27 motility is needed for motile cilia assembly *in vivo* (**Extended Data Fig. 8a, b**). Unlike *Kif27^−/−^*, *Kif27^R206K^* homozygous animals were viable with no hydrocephalus or nasal mucus accumulation (**Fig. 4a, b**). Motile cilia from the nasal epithelium and brain ventricles of *Kif27^R206K^*homozygous mice exhibited comparable morphologies and density as those observed for age-matched wt animals (**Fig. 4c**). Using wide-field live imaging, we evaluated cilia motility using respiratory epithelial tissues. Cilia beating frequencies were significantly reduced in *Kif27^−/−^* mice (**Fig. 4d; Supplementary Movies 1, 2**) but not in *Kif27^R206K^* homozygous animals (**Fig. 4e and Extended Data Fig. 8c; Supplementary Movies 3, 4**). Using expansion microscopy analysis, we found that SPAG6 and CP microtubules were present in more than 66% of respiratory ciliated cells from the *Kif27^R206K^* homozygous animals (**Fig. 4f, g and Extended Data Fig. 8d**). Cilia with abnormal structures were present, but not to the extent observed in *Kif27^−/−^* mice (**Fig. 4f**). The enhanced cilia beating frequency and restored SPAG+ CP+ motile cilia compared to *Kif27^−/−^* mice may be sufficient to revert PCD-related pathology in *Kif27^R206K^* homozygous animals. In mouse tracheal epithelial cells (mTECs) derived from *Kif27^GFP^* and *Kif27^R206K^* homozygous animals, KIF27^R206K^-GFP protein was made and expressed at a comparable level as KIF27-GFP (**Extended Data Fig. 8e)**. KIF27^R206K^-GFP co-localized with CP110 at the distal centrioles at the deuterosome stage and with CEP164 at the transition zone, but was lost from the tip of nascent cilia (**Fig. 4h-j**). The data is consistent with a scaffold function for KIF27 in motile cilia and suggest that the ATP-dependent motility of KIF27 may contribute to cilia tip targeting of the motor protein but is not necessary for KIF27 function in motile cilia biogenesis.

**Figure 4.**
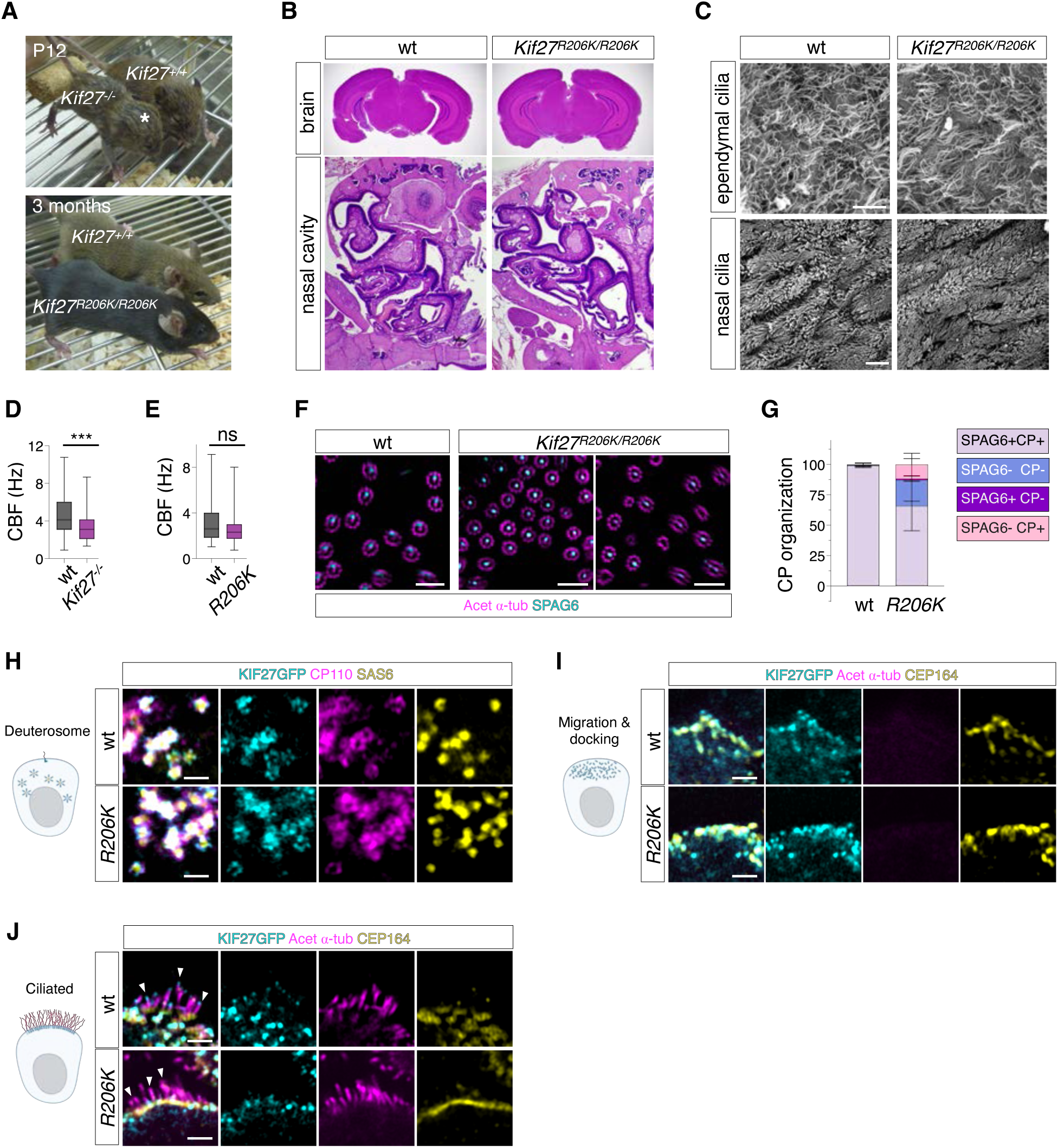
KIF27 motility is dispensable for PCD-related cilia functions. **(A)** At p12, *Kif27^−/−^* showed pronounced hydrocephalus compared to littermate wt (*Kif27^+/+^*). Asterisk indicates enlarged forebrain of the *Kif27^−/−^* mouse. At 3 months, *Kif27^R206K/R206K^* homozygous mice showed no difference in body growth or head shape compared to aged matched wt animals (*Kif27^+/+^*). All *Kif27^R206K/R206K^*mice were viable (n = 20). (**B**) Histology staining of the brain and nasal cavities from age-matched 4-month-old wt and *Kif27^R206K/R206K^* mice. **(C)** Scanning electron microscopy images of nasal epithelium and brain ependyma from age-matched 4-month-old wt and *Kif27^R206K/R206K^* mice. Scale bar = 10 μm. **(D and E)** The beating frequency significantly decreased in *Kif27^−/−^* (3.347 ± 1.546 Hz; n = 60, N = 2) compared to wt (4.492 ± 2.046 Hz; n = 60, N = 2); p = 0.008. Cilia beating frequencies were comparable in *Kif27^R206K/R206K^* animals (2.661 ± 1.336 Hz; n = 60, N = 2) compared to age-matching wt (3.154 ± 1.834 Hz; n = 60, N=2); p = 0.0956. The beating frequencies are shown as mean ± SD. **(F)** Tissue expansion images of wt and *Kif27^R206K/R206K^* nasal respiratory cilia showing the axonemal organization labeled with acetylated α-tubulin in magenta and the central pair SPAG6 signal in cyan for wt and *Kif27^R206K/R206K^*respiratory cilia. Scale bars = 4 µm. **(G)** Quantification of CP and SPAG6 organization in the wt (n = 300, N = 3) and *Kif27^R206K/R206K^* mice (n = 300, N = 3). **(H)** Endogenous KIF27-GFP and KIF27^R206K^-GFP (cyan) co-localize with distal centriole protein CP110 (magenta) during centriole amplification at the deuterosome stage in mTECs. SAS6 in yellow labels the centriole cartwheel. **(I)** KIF27-GFP and KIF27^R206K^-GFP (cyan) co-localize with transition fiber protein CEP164 (yellow) when the centrioles are docked. Centriolar microtubules are labeled by acetylated α-tubulin in magenta; the signal was faint at this stage. **(J)** KIF27-GFP (cyan) localizes at the tips of the short cilia but KIF27^R206K^-GFP (cyan) is absent from cilia tips. Cilia axonemes are labeled by labeled by acetylated α-tubulin in magenta. Cilia tips are indicated by arrowheads. Scale bars = 1 µm for **(H)**-**(J)**.

### KIF27 fine tunes the recruitment of ciliary transition zone proteins

CEP162 can promote TZ assembly in human retinal pigmented epithelial cells and is implicated in retina degeneration, a defect attributed to primary cilia dysfunction^58^. Given the association between KIF27 and CEP162, we hypothesized that the modes of action for KIF27 as a scaffold may take place at the TZ of motile cilia. To test this hypothesis, we first examined the sub-organellar architecture of the motile cilia TZ. Using lattice SIM super-resolution microscopy, we localized the TZ proteins CEP164, CEP290, MKS1, MKS3 and RPGRIP1L in respiratory ciliated epithelial cells from wt and *Kif27^−/−^*mice (**Fig. 5a**). In postnatal motile ciliated cells, the proximal-distal distances were measured for these markers relative to CEP164 to reflect the sub-organellar architecture of motile cilia TZ. The spatial organization for these ciliary TZ proteins in motile cilia was not identical to those observed for the primary cilium^59,60^ (**Fig. 5b**). In both wt and *Kif27^−/−^* cilia, these TZ proteins were organized in similar patterns in which MKS1 was the closest to CEP164, CEP290 and RPGRIP1L overlapped at a region distal to MSK1, and MKS3 was most distal to CEP164 (**Extended Data Fig. 9a**). Using conventional confocal microscopy, we then measured the fluorescence intensities for these TZ proteins in postnatal respiratory motile cilia. Signal intensities for CEP290, RPGRIP1L, and MKS1 were significantly reduced in *Kif27^−/−^* motile cilia, while MSK3 was unaffected (**Fig. 5c**). The data indicate KIF27 is required to recruit the correct amount of TZ proteins to motile cilia.

**Figure 5.**
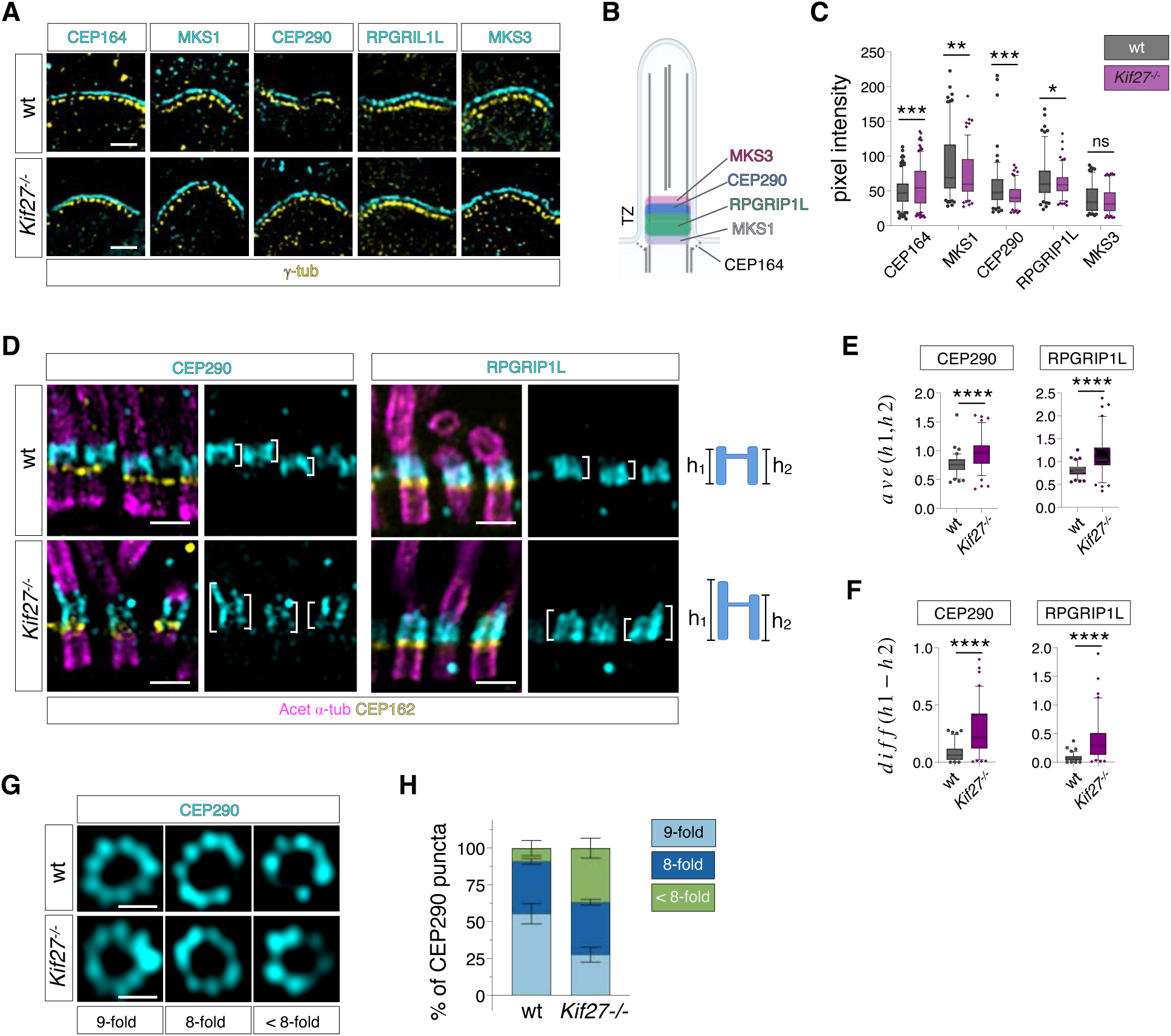
KIF27 is essential for maintaining motile cilia TZ integrity. **(A)** Super-resolution images of p17 mouse respiratory cilia transition fiber/transition zone markers CEP164, CEP290, MKS1 MKS3 and RPGRIP1L, all in cyan, with γ-tubulin (yellow) of the cilia for wt and *Kif27^−/−^* mice. Scale bar = 500 nm. **(B)** Schematic diagram of TZ organization in wt motile cilia. (**C**) Pixel intensities quantified using lattice SIM^2^ images of TZ markers for respiratory cilia from p17 wt and *Kif27^−/−^* mice. CEP164 (n for wt = 224, n for *Kif27^−/−^* = 231, N = 6 for each genotype, p-value < 0.0001, Student’s t-test), MKS1 (n for wt = 145, n for *Kif27^−/−^* = 137, N = 4 for each genotype, p-value = 0.0098, Student’s t-test), CEP290 (n for wt = 127, n for *Kif27^−/−^* = 120, N = 4 for each genotype, p-value = 0.0002, Student’s t-test), RPGRIP1L (n for wt = 138, n for *Kif27^−/−^* = 134, N = 4 for each genotype, p-value = 0.0301, Student’s t-test), MKS3 (n for wt = 134, n for *Kif27^−/−^* = 130, N = 4 for each genotype, p-value = 0.1522, Student’s t-test). **(D)** Expansion microscopy images of embryonic wt and *Kif27^−/−^* respiratory cilia stained with CEP290 and RPGRIP1L (cyan), acetylated ⍺-tubulin (magenta) and CEP162 (yellow). A representation of TZ organization is shown on the right for each genotype. Scale bar = 2 µm. (**E**) Average height (longitudinal axis) of CEP290 and RPGRIP1L domains measured with images taken using expansion microscopy shown in (**D**) (n = 90 for each genotype, N = 3). For RPGRIP1L, wt = 0.809 ± 0.133 μm vs *Kif27^−/−^* = 1.125 ± 0.392 μm, p < 0.0001; For CEP290, wt = 0.761 ± 0.157 μm vs *Kif27^−/−^*= 0.967 ± 0.263 μm, p < 0.0001. (**F**) Hight differences (longitudinal axis) for CEP290 and RPGRIP1L signals on each side of individual TZ measured from images taken with expansion microscopy shown in (**D**) (n = 90 for each genotype, N = 3). For RPGRIP1L, wt = 0.077 ± 0.064 μm vs *Kif27^−/−^* = 0.386 ± 0.356 μm, p < 0.0001; For CEP290, wt = 0.079 ± 0.069 μm vs *Kif27^−/−^*= 0.274 ± 0.299 μm, p < 0.0001. Plots show 10% - 90%. **(G)** *En face* images of CEP290 (cyan) for embryonic wt and *Kif27^−/−^* respiratory cilia. Scale bar = 1 µm. **(H)** Quantification of CEP290 puncta for individual transition zones (n = 90, N = 3 for each genotype). Error bars represent SD.

To investigate how KIF27 regulates TZ organization in greater detail, we used expansion microscopy to resolve the morphological nuances of TZ proteins. In ciliated respiratory cells isolated from wt mice, RPGRIP1L and CEP290 were enriched in a domain distal to CEP162 located at the base of cilia (**Extended Data Fig. 9b)**. From a lateral view, RPGRIP1L and CEP290 occupied a broader space than CEP164 and CEP162 (**Fig. 5d and Extended Data Fig 9b**). In *Kif27^−/−^* cilia, RPGRIP1L and CEP290 domains were distally expanded compared to wt and exhibited asymmetrical distribution along the axoneme (**Fig. 5d-f and Extended Data Fig. 9b**). TZ markers in motile cilia displayed a ring-like morphology comprised of discrete puncta with a 9-fold symmetry, similar to TZ markers in primary cilia^61^ (**Fig. 5g and Extended Data Fig. 9c)**. In *Kif27^−/−^* motile cilia, gaps between puncta were frequently present in TZ rings of CEP162 and RPGRIP1L (**Fig. 5g and Extended Data Fig. 9c**). The CEP290 TZ rings in *Kif27^−/−^* cilia often displayed < 9 puncta, and >60% of the rings showed 8 or less-than 8 puncta (**Fig. 5h**). The data collectively suggest that KIF27 is required for the molecular integrity of the motile cilium TZ.

### Proper barrier functions for motile cilia transition zone depend on KIF27

We hypothesized that an abnormally organized ciliary TZ may affect the ciliary recruitment of motility proteins, which may account for the structural and beating defects of motile cilia observed in *Kif27^−/−^* mutants. To test this model, we first defined the motile cilia proteome using mass spectrometry. Cilia fractions from mTECs derived from wt and *Kif27^−/−^* respiratory epithelia were isolated using a high-salt induced deciliation protocol (**Fig. 6a**). The protein abundance was obtained from three biological replicates for each genotype, and the abundancy ratio (WT/*Kif27 KO*) was normalized with an outer dynein arm component DNAH5, given that dynein outer arms were present in most *Kif27^−/−^*cilia based on TEM analysis **(Fig. 1e)**. Proteomics from three replicates consistently showed that essentially all components of the central pair apparatus were severely depleted from the *Kif27^−/−^*cilia fraction (**Fig. 6b**). Unexpectedly, the outer doublets associated MIPs, including the Tektin and PIERCE complexes^16^, were also significantly reduced in the *Kif27^−/−^* ciliary compartment (**Fig. 6b)**. The abundance of dynein arms and radial spoke proteins were not severely perturbed in mutant cilia. In contrast, core components of IFT components and the Bardet-Biedl Syndrome protein complex (BBSome) were mildly upregulated in the *Kif27^−/−^* cilia fraction. Conserved signaling modules, membrane proteins, and microtubule associated proteins were differentially affected by the lack of KIF27 (**Fig. 6b)**. In whole cell lysates collected from wt and *Kif27^−/−^* mTEC, comparable protein expressions for subunits of dynein arm complex, central pair apparatus, MIPs, and IFT complex were detected (**Extended Data Fig. 9d**), indicating that these motility proteins were not degraded. The data suggest that KIF27 is needed to promote the barrier function for the motile cilia TZ, and that TZ dysfunction may account for the missing structural proteins and impaired cilia motility observed in *Kif27^−/−^* cilia.

**Figure 6.**
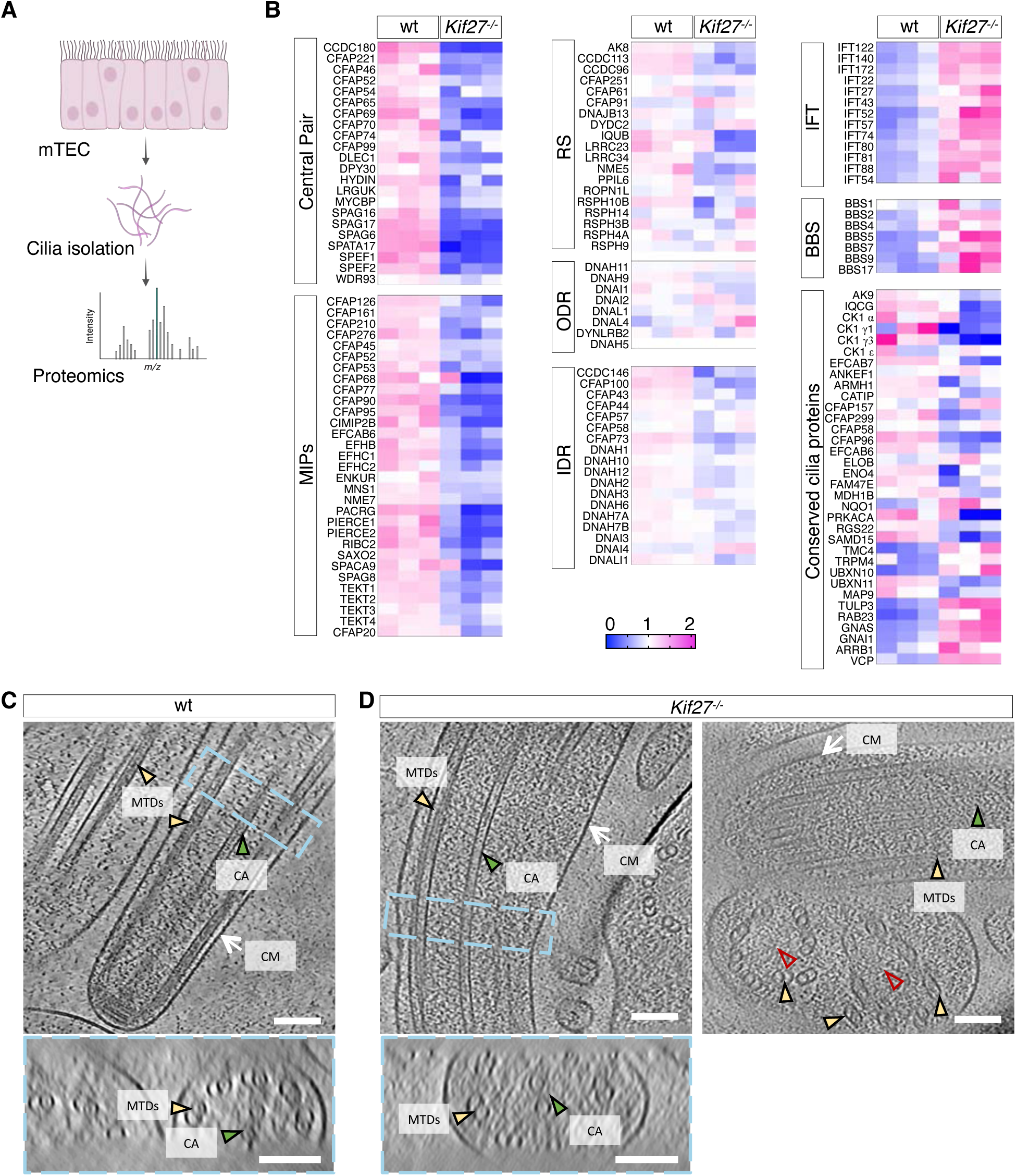
KIF27 is indispensable for proper barrier functions for motile cilia transition zone. **(A)** Schematic representation of proteomic analysis following cilia isolation. **(B)** Heat map of abundance levels normalized for dynein axonemal heavy chain 5 (DNAH5) in cilia isolated from wt and *Kif27^−/−^* mTEC (three replicates for each genotype) for representative proteins. Colors indicate fold change. MIPs: microtubule inner proteins; RS: radial spoke proteins; ODR: outer dynein complex; IDR: inner dynein complex. (**C-D**) Representative tomographic slices of cilia in wt (**C**) and *Kif27^−/−^* mTECs (**D**). In wt cilia, the CP microtubules are indicated by green arrowheads. In *Kif27^−/−^* samples, both CP+ and CP− cilia are present (red arrowheads indicate CP− cilia); CP− axonemes show a decrease in MTD numbers (from 9 to 8) or abnormally arranged MTDs. Compared to the wt, *Kif27^−/−^* cilia exhibit a significant amount of disordered densities in the ciliary compartment. Top: longitudinal central slices; Bottom: cross sections. CM: ciliary membrane, MTDs: microtubule doublets, CA: central pair apparatus. Scale bar = 100 nm.

To examine the spatial morphology of motile cilia axonemes in more detail, we employed *in situ* cryo-electron tomography (Cryo-ET) to reveal the native arrangements of axonemal microtubules and associated structural proteins. Vitrified motile ciliated cells dissociated from wt and *Kif27^−/−^*mTECs were first identified based on the expression of Centrin-GFP with correlative light and electron microscopy. These cells were thinned with a focused ion beam (FIB) and imaged by cryo-ET. Under the native cellular environment, microtubule outer doublets, CP microtubules, and CP apparatus were clearly visible in wt cilia (**Fig. 6c**). In *Kif27^−/−^* motile cilia, both CP+ and CP− cilia were observed, consistent with our TEM and expansion microscopy analysis (**Fig. 6d**). In those *Kif27^−/−^* mutant cilia with CP microtubules, densities correlated with the CP apparatus proteins appeared less organized spatially compared to the wt cilia. Regardless of the presence of CP, a conspicuous number of disordered densities were present within the luminal space of *Kif27^−/−^* mutant cilia and masked the densities of axonemal structural components. The extra density only appeared in *Kif27^−/−^* mutant cilia, which indicates that it is not part of the normal axonemal assembly and is consistent with ectopic retentions of proteins in the mutant cilia observed from cilia proteome. Collectively, the data suggest that KIF27-mediated TZ function is essential for the proper assembly of motile cilia by promoting the recruitment and correct incorporation of motility proteins into the ciliary compartment.

## Discussion

Despite the vital functions motile cilia perform, how mammalian motile cilia are built is not understood. In search of novel regulators of motile cilia biogenesis, we carried out an *in-silico* curation and identified KIF27 as a key candidate. Our study shows that mammalian KIF27 is essential for motile cilia biogenesis by controlling the correct molecular composition of the ciliary transition zone and suggest that KIF27 may be a new candidate gene for PCD. Although KIF27 is an ATP-dependent motile kinesin, it does not use its motility for cilia biogenesis. Instead, our data suggest an assembly pathway in which a KIF27-mediated scaffold at the transition zone is required to recruit structural proteins into mammalian motile cilia to drive motility. Since KIF27 homologues are conserved in most ciliated eukaryotes and absent from species without motile cilia^62^, KIF27 activities at the transition zone may define a unifying strategy to regulate motile cilia assembly in diverse species and cell types.

Motile cilia of *Kif27^−/−^* mice exhibit structure anomalies and defective waveforms, and gradually become shorter. The unusual ciliary defects set KIF27 apart from other known motile cilia kinesins, such as KIF9^30^ and KIF19^63^. In accordance with a synthetic KIF27 truncated protein^64^, the *in vitro* velocity for KIF27 motor dimers was much slower than those reported for the Kinesin-4 motors or IFT Kinesin-II, which is responsible for anterograde cilia transport^65^. Of the two homologues of KIF27, neither COS2 nor KIF7 is motile. Instead, both are required to tether the CI/GLI transcription factors to cytoplasmic and ciliary microtubules in *Drosophila*^66^ and mammals^35^, respectively, to mediate the graded response of Hh signaling. A parsimonious explanation is that, despite exhibiting ATP-dependent motility, a non-motile scaffold function may be selected for and conserved in the KIF27/KIF7/COS2 kinesin lineage in cilia-dependent and independent contexts. We show that mouse mutants homozygous for the *Kif27^R206K^*variant, which blocks ATP-dependent motility of KIF27, are alive without hydrocephalus or mucus accumulation in the nasal cavity, suggesting that a non-motile KIF27 is sufficient to drive motile cilia biogenesis in mammals. Nonetheless, a small fraction of abnormal axonemes is still present in *Kif27^R206K^* homozygous respiratory cilia. This incomplete rescue in cilia structure may be explained by a 40% decrease in *in vitro* microtubule binding duration observed for the KIF27^R206K^ motor dimers. In addition, KIF27^R206K^ failed to localize to the plus-ends of ciliary axonemes *in vivo*, suggesting that the ATP-dependent motility may account for the cilia tip tracking behavior of KIF27. Given the dynamic localizations for KIF27 in short and long cilia during ciliogenesis, the ATPase activity of KIF27 may contribute to the maintenance and stabilization of mature cilia. *In vitro* dynamic assays will clarify whether KIF27 can differentially regulate growth and stability of short and long microtubules, a combination of motor properties observed in some members of the Kinesin-4 family^36,46,47,52^.

In motile ciliated cells, KIF27 localizes to the transition zone in complex with CEP162, which crosslinks KIF27 with CEP290, a central scaffold of the MKS complex. KIF27-CEP162 association is mediated through the C-terminal cargo binding domain in KIF27 and the N-terminal non-coiled coil region of CEP162; the resulting complex creates a *de novo* microtubule binding interface which may contribute to microtubule stabilization at the ciliary transition zone. CEP290 can bind to microtubules and membrane lipids through two separate regions^67,68^. Ablation of either CEP290 domain is sufficient to disrupt ciliogenesis and induce mis-localization of membrane-bound ciliary cargos, including opsin and rhodopsin of the photoreceptors^68^. The proximity between KIF27 and CEP290 suggests a model in which the KIF27-CEP162-CEP290 scaffold is formed in nascent cilia, and this KIF27-mediated scaffold organizes and stabilizes the CEP290 complex at the transition zone to ensure a proper barrier function for cilia assembly and motility.

A connection between individual TZ proteins and cilia motility has been implicated in several species. Basalin, a *Trypanosoma brucei* specific TZ protein, is required for CP biogenesis by forming the basal plate, an electron dense structure proposed to nucleate and stabilize CP microtubules^69,70^. NPHP1 and NPHP4 are required for cilia structure and axonemal organization in the connecting cilia of photoreceptor cells and sperm tail^71–73^. Leber congenital amaurosis (LCA), a rare inherited form of early-onset eye diseases, causes severe visual impairment or blindness. A cohort of LCA patients with CEP290 variants were examined for the motile cilia morphologies based on their PCD-like symptoms, including chronic rhinitis and recurrent bronchitis. Nasal epithelial cells from these patients show axonemal defects, including missing CP, <9 microtubule doublets, as well as missing dynein arms^74^. Consistent with these findings, axonemal defects were reported in brain ependymal motile cilia in a *Cep290* knockout mouse model^75^.

How does the transition zone contribute to motile cilia assembly and function? Given that most TZ proteins have extensive coiled-coils, a structural feature that can mediate protein-protein interactions, one scenario is that the TZ is a size-exclusion barrier that can selectively guide the motility machineries to specific loading docks for their incorporation into the growing axoneme. In this model, TZ functions similarly in primary cilia and motile cilia to regulate ciliogenesis, during which the TZ barrier function is critical for IFT-mediated trafficking as well as cargo sorting and retention within the ciliary compartment^11,76^. Consistent with this idea, TZ dysfunction may account for the depletion of selected ciliary cargos, such as MIPs, and ciliary retentions of IFT and BBsome subunits observed in the motile ciliome of *Kif27^−/−^* mice and reported for *Cep290 Chlamydomonas* mutants^77,78^. Careful examination of the assembly dynamics of CP apparatus and MIPs during ciliogenesis will be necessary to clarify whether they are bona fide IFT cargos. Heterologous KIF27 expressed in HEK293T cells can bind to STK36^79^, a predicted serine-threonine kinase implicated in rare cases of PCD^80^. It remains an open question whether STK36 is tethered to the TZ through KIF27 to regulate phosphorylation-dependent barrier assembly and permeability, a phenomenon well characterized for other sorting machineries, such as the nuclear pore complex^81^, axon initial segment^82^, and septins^83,84^. An alternative and not mutually exclusive explanation is that KIF27, perhaps together with other TZ proteins, can directly interact with different sets of structural proteins to regulate their ciliary incorporation during ciliogenesis. Identification of the KIF27 interactome followed by functional validations will be required to shed light on the mechanistic details of KIF27-mediated TZ scaffold during motile cilia assembly.

The molecular identities of the TZ proteins and their functional interactions have been extensively characterized within the context of primary cilia formation and function, with an emphasis on TZ-mediated ciliary membrane protein trafficking^12^. Pathogenic variants have been identified for most genes encoding TZ proteins, underscoring the crucial role of the ciliary transition zone in embryonic development and tissue homeostasis. However, the phenotypes and severity of TZ-related ciliopathies vary widely across TZ genes. The lack of a clear genotype-phenotype correlation suggests that the composition and function of the TZ are not uniform, but may exhibit heterogeneity across different cell types and organs. The unexpected connection between a KIF27-mediated transition zone scaffold and the recruitment of motile cilia structural proteins highlights the gaps in our understanding of how the transition zone operates to regulate the assembly and function of mammalian motile cilia. Given the ubiquitous expression of TZ proteins in mammalian cells, it is likely that pathological variants of TZ genes can affect both primary and motile cilia and therefore, the manifestations of diseases associated with dysfunctional motile cilia may have been underestimated in ciliopathy patients. A mechanistic coupling between TZ dysfunction and motile ciliopathy represents an exciting area for future studies which may pave the way for expanding on the current list of motile ciliopathy genes and identifying new therapeutic strategies aimed at restoring cilia motility.

## Acknowledgement

We thank R. Subramanian, J. Hughes, B. Myers, and R. Fässler for valuable discussion and feedback on our work. We thank X. Duan, Y. Xia, X, Huang and S. Nakieln for comments on the manuscript. We thank P.T Chung (University of California, San Francisco) for providing the mouse Kif27-GFP construct. We thank T. Yang for sharing a protocol on expansion microscopy. We thank C.J. Khoo for technical support on protein purification. Imaging equipment are maintained by Centre for PanorOmic Sciences at the Faculty of Medicine, the University of Hong Kong. This work is supported by General Research Fund (M. He) and Canadian Institutes of Health Research (C. Hui).

## Author contributions

M.H and C.H conceptualized the project. H.O.C, S.M, Y.Y and C.H designed and established *Kif27^+/−^* and *Kif27^GFP^* mice. H.O.C, M.C, H.P and M.H performed and analyzed mouse experiments. G.L, P.L, and M.H designed and engineered *Kif27^R206K^* mice. M.C performed and analyzed imaging experiments. H.P made antibodies, purified proteins, performed and analyzed biochemical experiments. H.Q, H.P, G,F and S.T purified proteins, performed *in vitro* single molecule assays and analyzed results. H. P, Z. Liu and X. L performed and analyzed proteomics analysis. M.C, Q. W and Z.L performed and analyzed super-resolution imaging experiment. S.S.G made antibodies. W.W performed TEM experiments. M.H wrote the whole manuscript with assistance from H.P and M.C. All authors provided input to the manuscript.

## Materials and methods

### EXPERIMENTAL MODEL AND SUBJECT DETAILS

#### Mouse strains

All experimental procedures involving mice were performed in accordance with approved protocols by the Committee on the Use of Live Animals in Teaching and Research (CULATR) at the University of Hong Kong.

A loss-of-function mutation in *Kif27* (the *Kif27^−/−^* allele) was generated by replacing part of exon 1 and intron 1 with an internal ribosome entry site (IRES)-*lacZ* reporter gene and a PGK-*neo* cassette by homologous recombination in R1 ES cells. Correctly targeted ES cell clones were identified by Southern blot analysis and used to generate germline chimera. To delete the PGK-*neo* cassette at the targeted locus, *Kif27*^tm1/+^ mice were crossed with *NLS-*Cre transgenic mice, which show ubiquitous Cre recombinase expression (a gift from C. Lobe). Genotyping primers are:

P1-5’-AATCTGTTCAAAATGGCTCTG,

P2-5’-AGCCAAGTAAACAGACCAAC,

P3-5’- GGAGGGATCTTGGCCATGGTAAGCTG

P4-5’- CTGGCGACTGGATAGTTGTGGAAAGAGTC

To make a targeting plasmid to generate the *Kif27^GFP^*allele, three DNA fragments containing 3’ region of *Kif27* fused to *EGFP* gene were prepared for the left arm, and one fragment was prepared for the right arm. These three 4.4 kbp, 970 bp and 423 bp fragments were inserted into NotI site of pGKneo targeting vector as a left arm, which contains a *neomycin* cassette flanked with *loxP* sites. For right arm, a 3.3 kbp DNA fragment containing engineered SalI and EcoRV sites was obtained by PCR. This fragment was inserted into the targeting vector digested with SalI and EcoRV as a right arm. The targeting construct was electroporated into R1 ES cells and 96 neomycin-resistant colonies were screened by Southern blot analysis. Twenty correctly targeted clones were identified, and two of them were used to generate germline chimera. Genotyping primers are:

Kif27GFPcommonfrd: GTGTCGCAGAGAATTACGTC

Kif27GFPRvs: CAGATGAACTTCAGGGTCAGCT

Kif27GFPWTRvs: GAAGTTTTATTGTTCAGTGCC

To make the *Kif27^R206K^*allele, zygotes were collected from natural mating *Kif27^GFP/GFP^*mice. Reaction mix was prepared by including 20 ug Cas9 protein (Invitrogen), and 4.5ug guide RNA GAAAATAGCGTGGGATCTAC (Synthego) into Opti-MEM medium in total 60 μl. The mix was incubated at RT for 20 min to form Ribonucleoprotein (RNP) complex. Electroporation was carried out with Nepa21. In brief, 50 μl of the RNP complex was added into the chamber of the electrode. Zygotes were first washed with Opti-MEM for three times, followed by 1x wash with 10 μl RNP complex solution, and transferred to the chamber of the electrode. After electroporation, zygotes were taken out from the electrode and washed with fresh KSOM medium (Millipore) for three times and then cultured in KSOM media in the tissue culture incubator. The 2-cell stage embryos were transferred into the oviduct of CD1 (Charles River, France) pseudo-pregnant females (20 embryos per female). PCR amplifications were performed directly on alkaline lysates of ear biopsies from pups. Products of PCR were sent for Sanger Sequencing. The genotypes were by analyzing the sequencing data by Synthego ICE analysis software online. Genotyping primers are:

Kif27Exon4Frd: TTTGGGTTGCTTAGCAAAGGGT

Kif27Exon4Rvs: TCAAAATCCGAGGAGGACGG

#### Mouse tracheal epithelial cell culture

For mouse tracheal epithelial cells (mTEC) culture, cells were isolated from trachea of at least 8 weeks-old mice, cultured and expanded following the published protocol^1^. In brief, cells were isolate and dissociated from tracheas using 0.15% pronase (10165921001, Roche) in Ham’s F12 (11765070, Gibco) with Antibiotic-Antimycotic (15240112, Gibco), and with DNAse1 (07469, STEMCELL Technologies). Dissociated cells were then seeded onto Matrigel coated T75 flasks (For 3 adult tracheas) with KSFM medium (17005042, Gibco) supplemented as previously reported. When confluent, cells were further passaged with 1:3 ratio. When the cells reach confluency, the cells were dissociated with 0.25% trypsin (25200072, Gibco) and seeded onto Matrigel coated 0.33cm^2^ transwell insert (0.4µm pore size, 37024, SPL Life Science) with at least 100,000 cells per insert with mTEC proliferation medium. After the cells reach confluency, the medium is changed to mTEC differentiation medium and changed to air-liquid interface.

#### Human cell culture

HEK293T and U2OS cells were maintained at 37°C, 5% CO_2_ in DMEM Glutamax (Invitrogen) supplemented with 10% fetal bovine serum, 100 U/ml penicillin, 100 U/ml streptomycin. Transient transfections of plasmids into HEK293T and U2OS cells were performed using Lipofectamine 2000 and Lipofectamine 3000 (Invitrogen), respectively. Cells were regularly tested for mycoplasma contamination.

### METHOD DETAILS

#### Antibodies used in this study

Refer to key resources table, Table S2 for details regarding the primary antibodies.

#### Cloning and plasmids

To generate GFP-tagged KIF27 truncation constructs, fragments of KIF27 from full-length human KIF27 plasmid (a gift from R. Subramanian, Mass General Hospital/Harvard University, USA) were cloned into a modified pCBh vector (a gift from S.C. Kwon, The University of Hong Kong, Hong Kong SAR, China) with C-terminal GFP tag using Phusion polymerase (NEB). The linearized vector was prepared using EcoRI and AgeI restriction enzymes (NEB) and ligated with target insert fragment by Assembly Mix (TransGen). HA-tagged human CEP162 constructs were from W.J. Wong (National Yangming Chiao Tung University, Taiwan), and GFP-tagged mouse KIF27 full-length plasmid was from P.T. Chuang (US San Francisco, USA).

#### Immunoprecipitation

Cells were lysed in lysis buffer (20 mM Tris-HCl, 150 mM NaCl, 1% IGEPAL CA-630, 2 mM EDTA, 5% glycerol, protease inhibitor, pH 8) on ice for 30 min. The lysate was centrifuged at 16,100 g for 10 min at 4°C. Pierce Protein A/G magnetic beads (Thermo Scientific) pre-equilibrated in the lysis buffer were incubated with the lysate for 30 min at 4°C with rotation. Supernatant (‘precleared lysate’ and ‘Input’) was collected by magnetic stand and incubated with Pierce Anti-HA magnetic beads (Thermo Scientific) overnight at 4°C with rotation. For IgG isotype control samples, precleared lysate was incubated with Mouse IgG Isotype Control (Invitrogen) overnight at 4°C with rotation, then incubated with Pierce Protein A/G magnetic beads for 1 hour at RT. After the supernatant was collected and removed, the beads were resuspended in 50 µL 1XSDS sample buffer (‘Beads’ and ‘IgG’). The Input, Beads, and IgG samples were run on SDS-PAGE followed by transfer to PVDF membranes. For immunoblotting, antibodies were diluted in TBST with 5% non-fat milk.

#### Protein expression and purification: anti-GFP nanobody

The plasmid was transformed into Rosetta 2 (DE3) *Escherichia coli* cells (Sigma-Aldrich) and expressed at 16 °C with 0.3 mM IPTG for 16-18 hours. Bacterial pellets were lysed in TES buffer (200 mM Tris-HCl, 0.5 mM EDTA, 500 mM sucrose, pH 8). After incubating on ice for 30 min, lysate was cleared by ultracentrifugation, and the supernatant was incubated with Ni-NTA agarose (Qiagen) equilibrated in Ni-NTA binding buffer (20 mM Tris-HCl, 150 mM NaCl, pH 8.0) for 30 min. The resin was washed with wash buffer I (20 mM Tris-HCl, 900 mM NaCl, pH 8.0) then wash buffer II (20 mM Tris-HCl, 150 mM NaCl, 10 mM Imidazole, pH 8.0) and protein was eluted with Ni elution buffer (20 mM Tris-HCl, 150 mM NaCl, 250 mM imidazole, pH 8.0). The eluate was concentrated using 10K MWCO centrifugal filter (EMD Millipore) and further purified by size exclusion chromatography (Superdex 75 1660) in gel-filtration buffer (20 mM Tris-HCl, 150 mM NaCl, 10% glycerol, pH 8). After concentration, protein was snap-frozen with liquid nitrogen and stored at –80°C.

#### Protein expression and purification: KIF27 constructs

A fragment of KIF27 (aa350-560) was determined by sequence alignments by the Clustal Omega program using mouse KIF27 (Unirprot: Q7M6Z4), human KIF27 (Uniprot: Q86VH2), and human KIF7 (Uniprot: Q2M1P5). GFP-tagged KIF27 motor dimer (KIF27_1-570_), KIF27^E160A^, KIF27^R206K^ and tag-free KIF27 (aa 350-560) were cloned into a pFastBac expression vector with a tobacco etch virus (TEV) cleavable 6xHis-tag and a SUMOstar tag and expressed in SF9-baculovirus expression system as previously described^2^. The proteins were purified as previously described with modified protocol^3,4^. Cell pellet was lysed by dounce homogenizer in lysis buffer (50 mM phosphate, 300 mM NaCl, 5% glycerol, 30 mM Imidazole, 0.1mM ATP, 1 mM β-mercaptoethanol, 0.15% Tween-20 and 0.5% IGEPAL CA-630, pH 8), and the lysate was clarified by centrifugation. The lysate was loaded onto HisTrap HP column (Cytiva) pre-equilibrated with the lysis buffer. TEV protease diluted in digestion buffer (50 mM phosphate, 300 mM NaCl, 5% glycerol, 30 mM Imidazole, 0.1 mM ATP, 1 mM β-mercaptoethanol, pH 8) was loaded to the column and the protein was eluted after 5 rounds of on-column TEV protease digestion. Peak protein fractions were pooled and further purified by size exclusion chromatography (Superdex 75 1660 or Superdex 200 1660) in gel-filtration buffer (50 mM HEPES, 300 mM KCl, 10% glycerol, 5 mM β-mercaptoethanol, 0.1 mM ATP, 1 mM MgCl_2_, pH 7.4). Protein was snap-frozen with liquid nitrogen and stored at –80°C.

#### Protein expression and purification: TUBB4B/TUBA1A heterodimers

Affinity tag-free wildtype TUBB4B/TUBA1A heterodimers in unlabeled, biotinylated and fluorophore-labeled forms were expressed and purified from insect cells as previously described.^2^

#### Total Internal Reflection Fluorescence (TIRF) microscopy

Preparation of flow chambers and GMPCPP-stabilized TUBB4B/A1A microtubules were carried out as described^2^. To test whether the KIF27 motor proteins have ATP-dependent motility, single molecule imaging was carried out as described^5^ using reaction mixture of 20 pM protein in 1X BRB80 with 2 mM ATP, 1 mM MgCl_2_ (4 mM MgCl_2_ for KIF27^E160A^ and KIF27^R206K^), 5% sucrose and 0.2 mg/mL κ-casein. Time-lapse imaging was performed in Metamorph software (version 7.10.4) and images were taken every 1 s for total duration of 300 s. In addition, microtubule gliding assay was performed using a similar assay except that KIF27 proteins were immobilized to slide chamber using anti-GFP antibody (Abcam) before adding reaction mixture containing GMPCPP-stabilized microtubules. Time-lapse imaging was performed in Metamorph software (version 7.10.4) and images were taken every 1 s for total duration of 420 s. All *in vitro* TIRF-based imaging assays were performed using iLAS3 ring-TIRF microscopy (Inverted: Nikon Ti2-E) with 100x 1.49 oil lens. The microscope lens was kept warm at 37 °C with objective heater, and excitation was achieved using 488 nm and 561 nm laser.

#### Histological and skeletal analysis

Histological analysis and skeletal analysis of two weeks old mice were performed using alcian blue/alizarin red bone staining as described^6^. Briefly, for brain and nasal section histology, whole tissue samples were fixed with 10% neutral buffered formalin overnight in RT. Then the whole tissue was dehydrated with and stored in 80% ethanol. For nasal samples, decalcification was done with 15% formic acid. Whole tissue samples were then paraffinembedded and sectioned. H&E staining was done using the sections.

#### Electron Microscopy

For scanning electron microscopy, trachea, nasal epithelium and brain tissue were isolated from mice of designed genotypes. Large airway, nasal epithelium and brain ventricles were collected. Tissue samples were fixed with 2% paraformaldehyde (15952-10S, Electron Microscopy Science) and 2.5% glutaraldehyde (16539-30, Electron Microscopy Science) in 0.1 M sodium cacodylate buffer for 24 hours at RT. The fixed tissue was washed three times, 10 min each with 0.1M sodium cacodylate buffer prepared with 0.4M sodium cacodylate buffer (11654, Electron Microscopy Science) diluted with filtered PBS and dehydrated in ethanol series (25%, 50%, 75% and 100%) (refer to Kif7 paper). Following dehydration, critical-point drying was done in Electron Microscopy Unit in the University of Hong Kong. Images were taken with Hitachi S34000N VP SEM.

For transmission electron microscopy, trachea tissues were collected from postnatal day 15 (p15) wild-type (wt) and *Kif27^−/−^*animals and fixed with 2% paraformaldehyde and 2.5% glutaraldehyde with 1% tannic acid in 0.1 M cacodylate buffer. The samples were then post-fixed in 1% OsO4 in 0.1 M cacodylate buffer at RT for 30 min, stained with 1% uranyl acetate at RT for 1 hour, dehydrated in a graded series of ethanol, infiltrated, and embedded in Spurr’s resin. Thin sections (80 nm) were stained with 4% uranyl acetate and Reynold’s lead citrate for 10 min and examined with an electron microscope (T FEG-TEM; FEI Tecnai G2 TF20 Super TWIN).

#### Immunofluorescence confocal microscopy

Samples were fixed using 2-4% paraformaldehyde (PFA) for 10 to 20 min at RT. For certain transition zone markers, samples were fixed with ice cold methanol for 20 min followed by acetone for 1 min. For tissue, after fixation and rinsing with 1xPBS, they were submerged 30% sucrose in PBS overnight and embedded in OCT tissue freezing medium (14020108926, Leica). Tissue cryo-blocks were sectioned to 10 to 15 μm slices attached to Superfrost Plus slides (J1800AMNZ, Thermo Fisher). For mTEC fixation, the whole insert was fixed for 10 min with 2% PFA in RT. For immunostaining, tissue sections or cells were blocked for 1 hour at RT in blocking buffer (1xPBS + 1% heat inactivated serum (Jackson ImmunoResearch) + 0.3% Triton X-100 (Sigma)) and incubated with the primary antibodies overnight at 4°C. Cells were stained with secondary antibodies for 1-2 hours at RT. Samples were imaged on Zeiss LSM 980 inverted confocal microscope with 40X (1.4 NA) oil objective at the Centre for PanorOmic Sciences (CPOS) of the University of Hong Kong Faculty of Medicine.

#### Lattice SIM super-resolution microscopy

Imaging samples were prepared using the same protocol for regular confocal microscopy. Lattice SIM super-resolution data collection and analysis. Lattice SIM datasets were collected on a Carl Zeiss ELYRA 7 microscope at Biosciences Central Research Facility, Hong Kong University of Science and Technology. In brief, the fluorophores were excited with 488 nm (500 mW, tuned to 0.5-10% with the ZEN software) and 561 nm (500 mW, tuned to 0.5-10% with the ZEN software) lasers and collected with a 63x/1.46 oil immersion objective lens with a 1.6x extra magnification lens inserted in the light path. The collected photons were split into two halves by a beam splitter, and the fluorescence was filtered by bandpass mirrors (495-550 for the green channel and 570-640 for the red channel respectively) before reaching the cameras. Two PCO.edge 4.2 sCMOS cameras were used to collect images from two channels simultaneously. Exposure time and laser power were optimized to reach higher image contrast. For each field of view, the lattice illumination pattern was applied on the focal plane and 13 phases were collected for image reconstruction. A 5-10 μm Z stack was collected for each dataset with a thickness of 100 nm per Z step.

#### Tissue ultrastructure expansion microscopy

Fixed cells or tissue section samples were incubated with infusion buffer (0.7% PFA and 1% Acrylamide in PBS) for 5h in 37°C. Samples were then washed with 1XPBS 3 times, 10 min each. Gelation chamber was made by placing two #1.5 square cover glasses on each side of the samples. Gelation solution (19% sodium acrylate, 10% acrylamide, 0.1% BIS, 0.5% TEMED, 0.5% APS in 1X PBS) was directly placed on top of the samples and parafilm covered slide is placed on top of the sample slide with the gelation solution. For complete gelation, sample was incubated at 37°C for 1h. Sample embedded in the gel was carefully removed from the glass slide using a blade and incubated in denaturation buffer (200mM SDS, 200mM NaCl and 50mM Tris-HCL dissolved in H_2_O) for 90 min at 95°C. After denaturation, the gel was washed twice with 1X PBS and expanded overnight with ddH_2_O exchange at least three times. For immunostaining, gel was washed with 1X PBS three times for 30 min in total. Gel was incubated with primary antibodies in IF buffer overnight on shaker in RT and washed three times with IF buffer 10 min each before secondary antibody incubation. Secondary antibody and DAPI incubation were also done overnight on shaker in RT and the gel was washed three times with IF buffer and expanded with H2O with three exchanges for at least 30 min after each exchange. Expanded gels were placed on poly-L-Lysine (500µg/ml) coated glass bottom dish (P35G-1.5-20-C, Mattek) for imaging. Images were acquired using Zeiss LSM 980 inverted confocal microscope with 20X (0.8 NA), 40X (1.4 NA) oil, and 63X (1.2 NA) water immersion lenses at CPOS of the University of Hong Kong.

The reagents used for ultrastructure expansion microscopy are: sodium acrylate (BD151354 Bide Pharmatech), acrylamide (A9099-100G, Sigma), N, N-methylenbisacrylamide (BIS), tetramethylethylendiamine (TEMED, 15524010, Invitrogen), ammonium persulfate (APS, HC2005, Thermo Fisher), and SDS (1610301, Bio-Rad).

#### Cilia live-imaging

Large airways were first isolated from postnatal or adult mice. Tracheas were then cut longitudinally to expose the lumen and cut into a total of 6-8 pieces per trachea. Trachea pieces were saved in PBS on ice until live-imaging and put to warmed DMEM with 10% FBS and Penicillin Streptomycin for imaging. The luminal side of the tracheal section faced the cover-slip of a 35mm MatTek dish. Live imaging was captured by Nikon Ti2E Widefield Florescence Microscope was used with 100x 1.45 oil lens. Temperature was set to 37°C using water chamber to maintain humidity. 16-bit Images were acquired with 30ms/frame.

#### Cilia isolation

Mature ALI cultures in 12-well size transwell were washed with 1xPBS three times then transwell membrane was transferred to pre-chilled tube containing deciliation buffer (10 mM Tris, 50 mM NaCl, 10 mM CaCl_2_, 1 mM EDTA, 0.1% Triton X-100, 7 mM β-mercaptoethanol, protease inhibitor, pH 7.5) and mixed vigorously. Cells were centrifuged at 1000 g for 1 min at 4°C to pellet debris. Supernatant was transferred to new pre-chilled tube on ice and centrifuged at 12,000 g for 5 min at 4°C to collect cilia pellet. The cilia pellet was resuspended in 100 µL resuspension buffer (30 mM HEPES, 25 mM NaCl, 5 mM MgSO_4_, 1 mM EGTA, 0.1 mM EDTA, 0.1 mM DTT, protease inhibitor, pH 7.3) and stored at −80°C until mass spectrometry

#### Mass spectrometry

The pellet was suspended in SDS lysis buffer (4% SDS, 150 mM NaCl, 25 mM HEPES, pH 7.5) with protease inhibitor and lysed by sonication (25% amplitude; 5 s on, 10 s off for 2 min). Proteins were diluted by lysis buffer (150 mM NaCl, 25 mM HEPES, pH 7.5) to a SDS concentration of 1%. Lysate was centrifuged at 21,000 g, 4 °C, for 15 min and the supernatant collected. The final lysate protein concentration was determined by BCA assay (Thermo Fisher Scientific). Then, the lysate was treated with DTT to achieve a final concentration of 10 mM. The mixture was then incubated at 50 °C for 30 min. After that, a final concentration of 30 mM iodoacetamide was added and incubated at RT for 1 hour in dark to alkylate the reduced cysteine residues. The cell lysate was precipitated by adding 4× volume of ice-cold acetone. The mixture was then incubated overnight at −20 °C. The next day, it was centrifuged for 10 min at 20,000 g and 4 °C to separate the precipitate from the supernatant. The resultant precipitates were washed once with ice-cold methanol and dried using a CentriVap Vacuum Concentrator (Labconco). Protein precipitates were resuspended in 50 mM TEAB buffer (Thermo Fisher Scientific) and digested with trypsin (Thermo Fisher Scientific) at 37 °C overnight at a mass ratio of 1:50. A second digestion was performed at 37 °C for 4 hour with additional trypsin at a mass ratio of 1:100. After digestion, peptides were then desalted by Sep-Pak Vac tC18 cartridges (Waters) and dried. Peptides were further fractionated with XBridge Peptide BEH C18 column (300 Å (30 nm), 3.5 μm, 2.1 mm × 250 mm) on an Agilent 1260 Infinity II system with a gradient of 2–80% acetonitrile in 10 mM NH_4_HCO_3_, pH 10.0 for 50min and dried using a vacuum concentrator. Peptides were resuspended in 0.5% formic acid for analysis. Samples were analyzed on a Q Exactive Plus mass spectrometer equipped with an Ultimate 300 RSLCnano system using PepMap 100 C18 trapping cartridge (5 μm, 300 μm × 5 mm) and PepMap RSLC C18 microcapillary column (2 μm, 75 μm × 25 cm) as the analytical column. Data were acquired using Thermo Xcalibur 4.3 software. Peptides were separated with a 4–40% acetonitrile gradient in 0.1% formic acid over 90 min at a flow rate of 300 nlmin^−1^. Electrospray ionization was performed by applying a voltage of 2,100 V through the PEEK junction at the inlet of the microcapillary column. MS analysis was performed in data-dependent mode. The survey scan was performed in the Orbitrap within the range 300– 1,800 *m*/*z* using the resolving power of 70,000, and then the 20 most intense peaks were selected for higher-energy C-trap dissociation (HCD) fragmentation using a precursor isolation width window of ±1.5 *m*/*z* and a resolution of 17,500. The automatic gain control for survey and MS2 scans was set to 1 × 10^6^ ions and 5 × 10^4^ ions, respectively. The maximum ion accumulation times for survey and MS2 scans were 30 ms and 50 ms, respectively. Normalized collision energy for HCD-MS2 was 28%. Ions with a single charge, more than eight charges or an unassigned charge state were excluded from MS2 analysis.

#### Cell vitrification and cryo-FIB milling

mTECs derived from *Kif27^+/+^; Cent-GFP* and *Kif27^−/−^;Cent-GFP* P15-P17 animals. After 4 weeks of ALI induced differentiation, cells were dissociated from the transwell inserts using TryplE (Thermo Fisher Scientific) prior to vitrification. Holey carbon-coated copper (R 2/1, 200 mesh) (Quantifoil) grids were glow discharged for 45 seconds at 15 mA in a PELCO easiGlow™ Glow Discharge Cleaning System. 4 µl of dissociated mTEC cells in DMEMF/12 (1500 cell/µl) was applied to the carbon side of glow-discharged grids, and 3 µl of cell culture medium to the back side in Leica GP2 (Leica Microsystems) chamber (37 °C, 95% humidity). After blotting for 7-8 seconds from the back side, grids were plunge-frozen in liquid ethane. The grids were stored in a liquid nitrogen tank until further use.

Grids were loaded to Thunder Imager EM CRYO CLEM (Leica Microsystems) to identify the GFP-positive target cells. The coordinates of GFP-positive cells were exported in Maps format and transferred to Aquilos 2. The grids were transferred to Aquilos 2 Cryo-FIB (Thermo Fisher Scientific) to prepare the cellular lamellae. The grids were firstly sputter coated for 30 s, followed by GIS coating for 20 s. The cells were automated thinned to ∼350 nm (ion beam current from 1 nA to 30 pA) using AutoTEM (Thermo Fisher Scientific) and manually polished to achieve a thickness around 200 nm.

#### Cryo-ET data collection and Tomogram reconstruction

Data collection of lamellae was performed by using a 300 kV Krios G4 Cryo-TEM (Thermo Fisher Scientific) equipped with a selectris energy filter with a nominal magnification of 53,000 (pixel size: 2.417 Å). After pre-tilt angle estimation of lamellae, tilt series were acquired from −64° to 48° at 2° increments with 2e^−^/Å^2^ per tilt using PACE-tomo in serialEM. The target defocus of tilt-series was set to −4 to −6 μm. The energy filter slit width was set to 20 eV and zero-loss peak was collected.

The raw EER movies were motion-corrected by RELION, with the application of 5 × 5 patches. Using IMOD, the dark images were removed by calculating the mean intensity of motion-corrected images using imod clip command. The motion-corrected images were then stacked into individual tilt series, which were aligned in Aretomo2, following the previously established method^7^. The tomograms were reconstructed at bin 2, followed by denoising procedures using cryoCARE^8^ with default parameters.

### QUANTIFICATION AND STATISTICAL ANALYSIS

#### Quantification for confocal microscopy images

Images were processed with Airyscan Processing using ZEN software and quantifications such as signal intensities were done using FIJI/ImageJ software.

#### Quantification for lattice SIM super-resolution images

The images were reconstructed with the ZEN software with the SIM square mode, which provides a lateral resolution of ∼ 60 nm. Channel alignment was performed with fiducial fluorescent beads. Image analysis such as pixel distance measurements were done using ZEN Blue Software.

#### Quantification for cilia live-imaging

Kymographs of 16-bit videos were generated using FIJI/ImageJ and total number of peaks (ciliary beat cycles) in 30 secs were counted.

#### Data analysis for proteomics

Thermo Proteome Discoverer (v.3.0 SP1) was used to search raw data against the Uniprot Mus musculus (Mouse) database (UP000000589). The following modifications were included: cysteine carbamidomethylation as fix modifications, methionine oxidation, protein N-terminus acetylation and methionine loss as variable modifications. Mass error tolerance was set as 20 p.p.m. for MS1 spectra and 0.06 Da for MS2 spectra, and maximum missed cleavages were set to 2. The FDR for peptide and protein was set to 0.01. P values were calculated using two-way Student’s t-test with three replicates and adjusted using the Benjamini–Hochberg procedure.

#### Data analysis for TIRF assays

Using Mosaic plugin function in ImageJ, the background of TIRF images was subtracted. Analysis of the velocity and binding time of track were performed in Ilastik (version 1.4.0) and MATLAB (version R2022a). The codes were acquired from Github as published (Github repository kjran/Kymo_analysis DOI:10.5281/zenodo.7699922). Kymographs pre-processed by Ilastik were further loaded to the Matlab. The graphs (Gaussian distribution) and statistical tests were performed using Graphpad Prism.

#### Statistical tests

Methods for statistical analysis and numbers of samples measured in this study are specified in figure legends. The error bars indicate standard deviation or standard error. Two-tailed Student’s t test were performed using Graphpad Prism software for data and are indicated in figure legends.

#### Key resources table

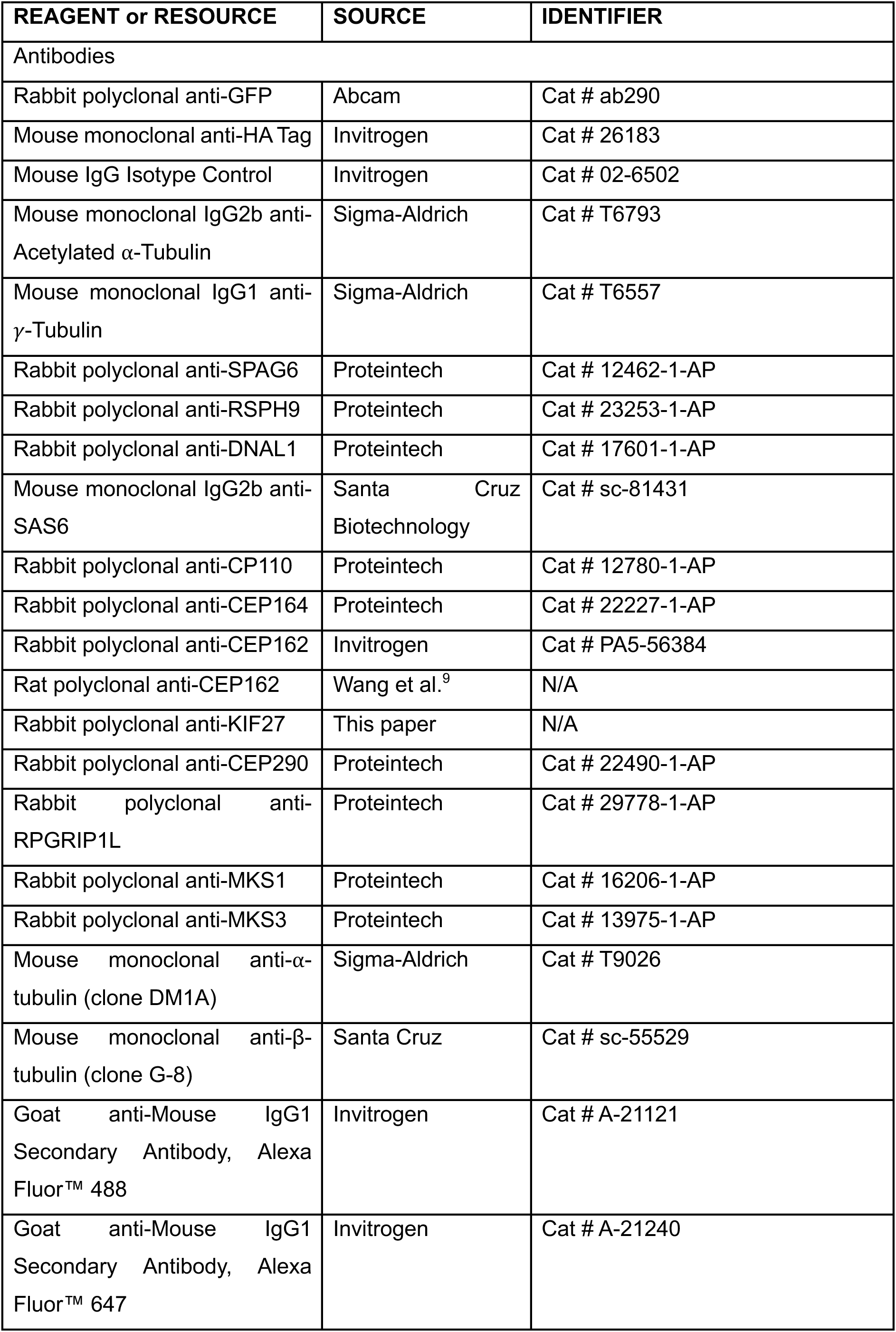
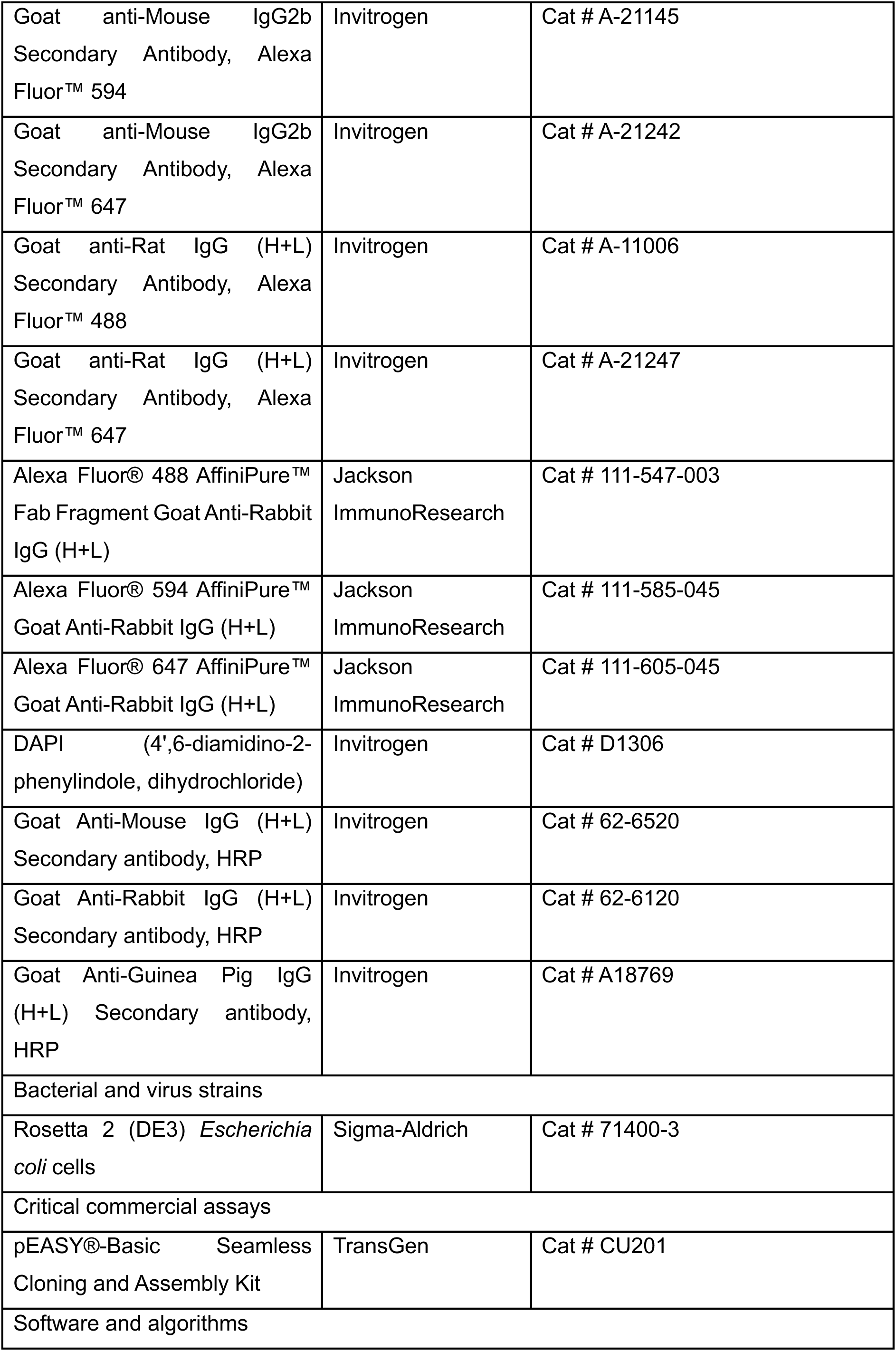
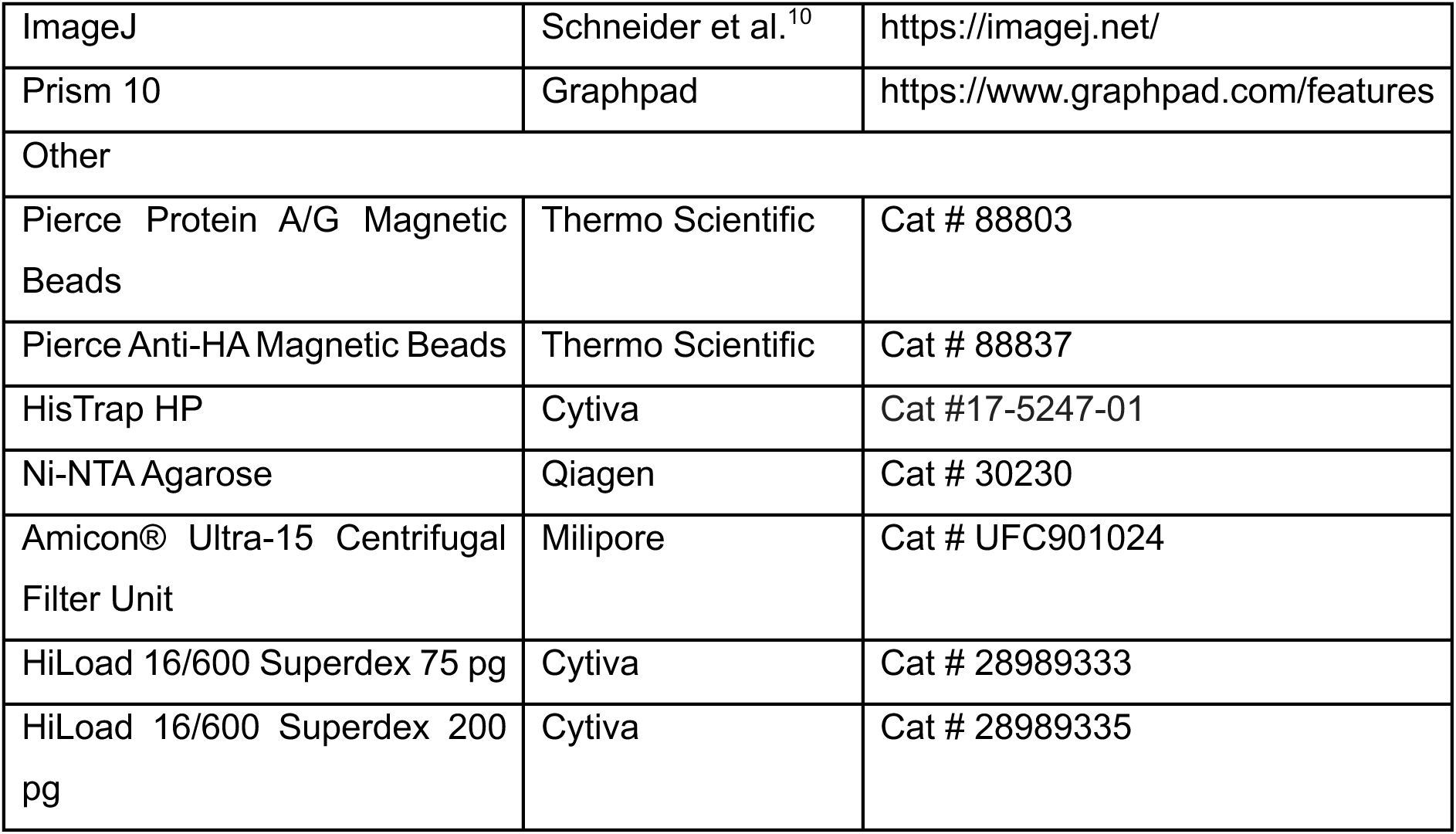

| REAGENT or RESOURCE | SOURCE | IDENTIFIER |
| --- | --- | --- |
| Antibodies |  |  |
| Rabbit polyclonal anti-GFP | Abcam | Cat # ab290 |
| Mouse monoclonal anti-HA Tag | Invitrogen | Cat # 26183 |
| Mouse IgG Isotype Control | Invitrogen | Cat # 02-6502 |
| Mouse monoclonal IgG2b anti-Acetylated α-Tubulin | Sigma-Aldrich | Cat # T6793 |
| Mouse monoclonal IgG1 anti-γ-Tubulin | Sigma-Aldrich | Cat # T6557 |
| Rabbit polyclonal anti-SPAG6 | Proteintech | Cat # 12462-1-AP |
| Rabbit polyclonal anti-RSPH9 | Proteintech | Cat # 23253-1-AP |
| Rabbit polyclonal anti-DNAL1 | Proteintech | Cat # 17601-1-AP |
| Mouse monoclonal IgG2b anti-SAS6 | Santa Cruz Biotechnology | Cat # sc-81431 |
| Rabbit polyclonal anti-CP110 | Proteintech | Cat # 12780-1-AP |
| Rabbit polyclonal anti-CEP164 | Proteintech | Cat # 22227-1-AP |
| Rabbit polyclonal anti-CEP162 | Invitrogen | Cat # PA5-56384 |
| Rat polyclonal anti-CEP162 | Wang et al.<sup>9</sup> | N/A |
| Rabbit polyclonal anti-KIF27 | This paper | N/A |
| Rabbit polyclonal anti-CEP290 | Proteintech | Cat # 22490-1-AP |
| Rabbit polyclonal anti-RPGRIP1L | Proteintech | Cat # 29778-1-AP |
| Rabbit polyclonal anti-MKS1 | Proteintech | Cat # 16206-1-AP |
| Rabbit polyclonal anti-MKS3 | Proteintech | Cat # 13975-1-AP |
| Mouse monoclonal anti-α-tubulin (clone DM1A) | Sigma-Aldrich | Cat # T9026 |
| Mouse monoclonal anti-β-tubulin (clone G-8) | Santa Cruz | Cat # sc-55529 |
| Goat anti-Mouse IgG1 Secondary Antibody, Alexa Fluor™ 488 | Invitrogen | Cat # A-21121 |
| Goat anti-Mouse IgG1 Secondary Antibody, Alexa Fluor™ 647 | Invitrogen | Cat # A-21240 |
| Goat anti-Mouse IgG2b Secondary Antibody, Alexa Fluor™ 594 | Invitrogen | Cat # A-21145 |
| Goat anti-Mouse IgG2b Secondary Antibody, Alexa Fluor™ 647 | Invitrogen | Cat # A-21242 |
| Goat anti-Rat IgG (H+L) Secondary Antibody, Alexa Fluor™ 488 | Invitrogen | Cat # A-11006 |
| Goat anti-Rat IgG (H+L) Secondary Antibody, Alexa Fluor™ 647 | Invitrogen | Cat # A-21247 |
| Alexa Fluor® 488 AffiniPure™ Fab Fragment Goat Anti-Rabbit IgG (H+L) | Jackson ImmunoResearch | Cat # 111-547-003 |
| Alexa Fluor® 594 AffiniPure™ Goat Anti-Rabbit IgG (H+L) | Jackson ImmunoResearch | Cat # 111-585-045 |
| Alexa Fluor® 647 AffiniPure™ Goat Anti-Rabbit IgG (H+L) | Jackson ImmunoResearch | Cat # 111-605-045 |
| DAPI (4',6-diamidino-2-phenylindole, dihydrochloride) | Invitrogen | Cat # D1306 |
| Goat Anti-Mouse IgG (H+L) Secondary antibody, HRP | Invitrogen | Cat # 62-6520 |
| Goat Anti-Rabbit IgG (H+L) Secondary antibody, HRP | Invitrogen | Cat # 62-6120 |
| Goat Anti-Guinea Pig IgG (H+L) Secondary antibody, HRP | Invitrogen | Cat # A18769 |
| Bacterial and virus strains |  |  |
| Rosetta 2 (DE3) *Escherichia coli* cells | Sigma-Aldrich | Cat # 71400-3 |
| Critical commercial assays |  |  |
| pEASY®-Basic Seamless Cloning and Assembly Kit | TransGen | Cat # CU201 |
| Software and algorithms |  |  |
| ImageJ | Schneider et al.<sup>10</sup> | https://imagej.net/ |
| Prism 10 | Graphpad | https://www.graphpad.com/features |
| Other |  |  |
| Pierce Protein A/G Magnetic Beads | Thermo Scientific | Cat # 88803 |
| Pierce Anti-HA Magnetic Beads | Thermo Scientific | Cat # 88837 |
| HisTrap HP | Cytiva | Cat #17-5247-01 |
| Ni-NTA Agarose | Qiagen | Cat # 30230 |
| Amicon® Ultra-15 Centrifugal Filter Unit | Milipore | Cat # UFC901024 |
| HiLoad 16/600 Superdex 75 pg | Cytiva | Cat # 28989333 |
| HiLoad 16/600 Superdex 200 pg | Cytiva | Cat # 28989335 |

---

**Supplementary Figure 1.**
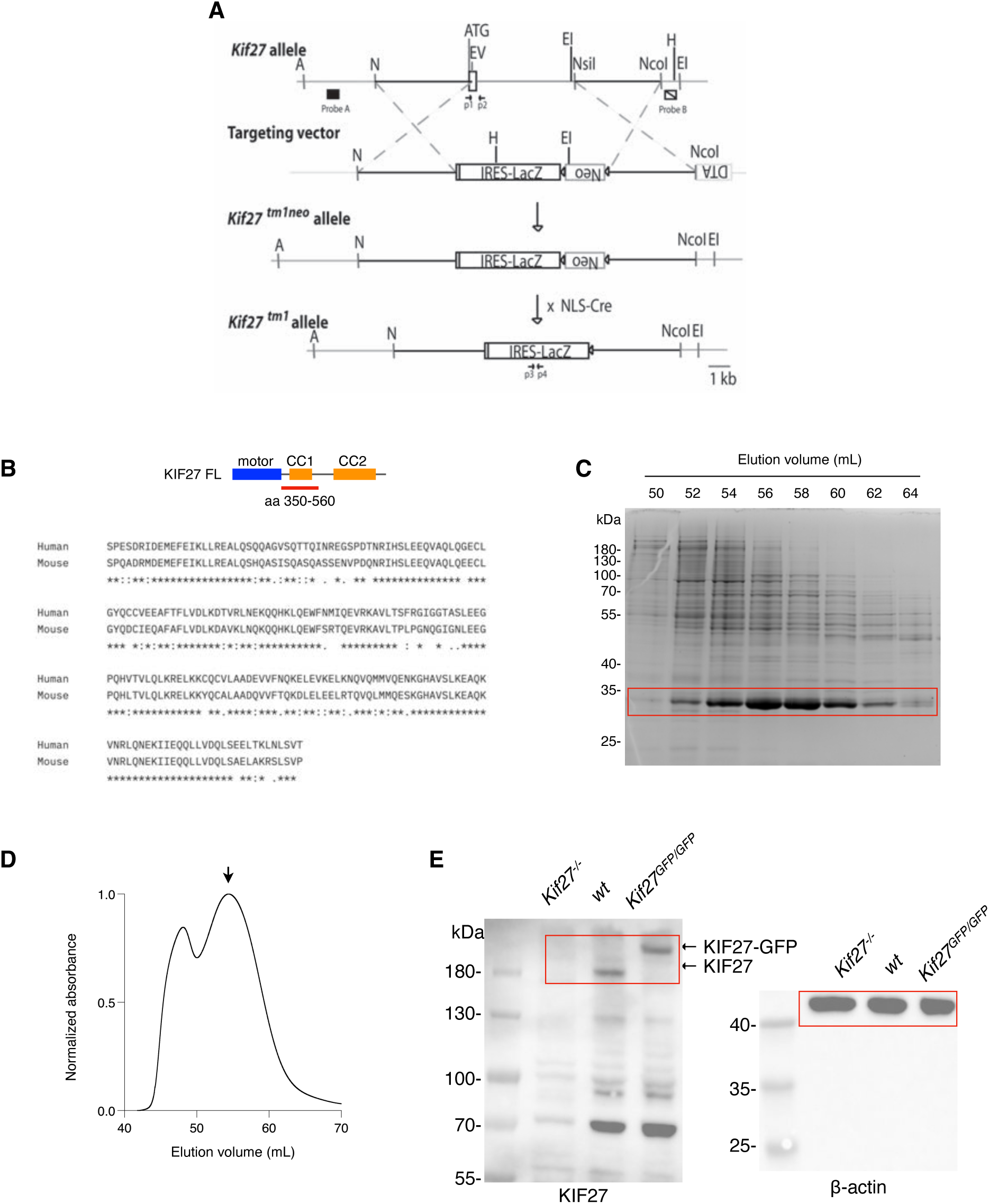
Generation of *Kif27^−/−^* mouse line and anti-KIF27 antibody. **(A)** Targeted mutation of the *Kif27* locus in ES cells was performed by inserting an IRES-*LacZ* gene (IRES-NLS-*LacZ*-PolyA) and the neomycin selection gene (PGK-neo) with stop codon into exon 1. Homologous recombination result in a change of size of ApaI fragment detected by the 5’probe from 17 kb (WT) into 9kb (Mut), while the EcoRI fragment detected by the 3’ probe shifts from 5kb (WT) into 6.6kb (Mut). A; ApaI, EI; EcoRI, EV; EcoRV, H; HindIII, N; NotI. **(B)** Domain organizations of full-length human KIF27_1-1401_ and region selected as the antigen (amino acid 350-560) indicated by red bar. ClustalW sequence alignment of human and mouse KIF27 for the selected region. **(C)** Coomassie blue-stained SDS-PAGE analysis of protein samples collected during gel filtration highlighted by red rectangle. **(D)** Elution profile from size-exclusion chromatography of KIF27 antigen (peak volume, 54.41 mL). The arrow indicates the expected position of purified protein. **(E)** Immunoblot for anti-KIF27 antibody verification using testis tissue lysates of *Kif27^−/−^*, wildtype (wt), *Kif27^GFP/GFP^*adult animals. β-actin served as loading control. Regions of interest are highlighted by red rectangles.

**Supplementary Figure 2.**
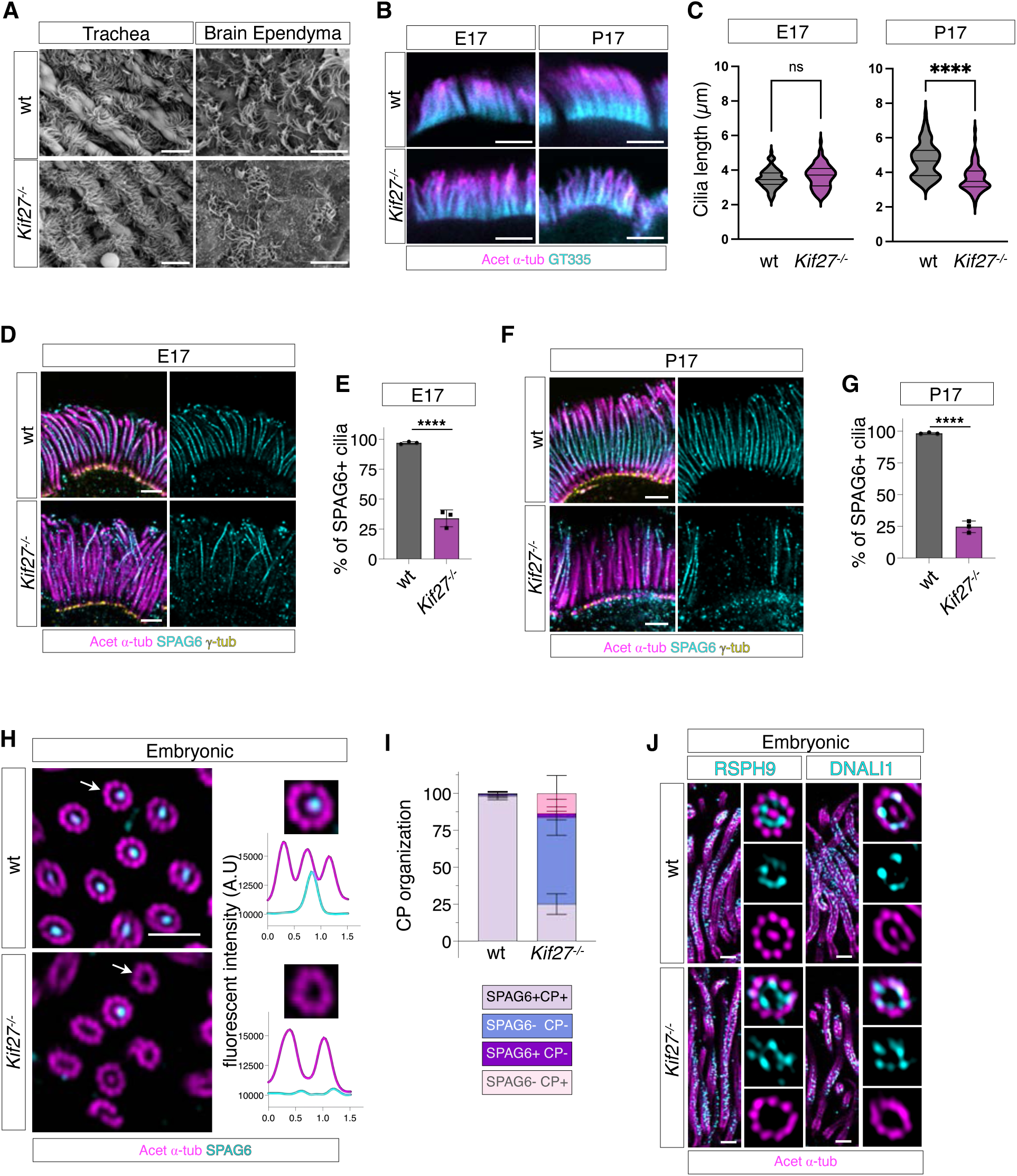
KIF27 is indispensable for proper motile ciliogenesis. **(A)** Scanning electron microscopy images of P20 wt and *Kif27^−/−^* mice trachea and brain ependyma. Scale bar = 10 µm. **(B)** Confocal immunofluorescence images of wt and *Kif27*^−/−^ nasal respiratory cilia stained with acetylated ⍺-tubulin (magenta) and polyglutamylated tubulin (GT335) (cyan) markers. Scale bar = 3 µm. **(C)** Quantification of cilia length based on the pixel distance of acetylated ⍺-tubulin for wt and *Kif27*^−/−^ mice respiratory airway for embryonic and postnatal stages (n=90, N=3 for each condition). Difference for cilia length in embryos was 0.1389 ± 0.09162 μm (p-value = 0.1313, Student’s t-test), and for postnatal it was 1.044 ± 0.1304 μm (p-value < 0.0001, Student’s t-test). **(D and F)** Sideview images of embryonic **(D)** and postnatal **(F)** mouse respiratory cells showing the missing SPAG6 signals in the *Kif27*^−/−^ mice using ultrastructure expansion microscopy. Scale bar = 2 µm. **(E and G)** Quantification of proportion of cilia with SPAG6 signals in cyan (n=300, N=3 for wt and *Kif27*^−/−^ at each developmental stages). Acetylated ⍺-tubulin in magenta labels the axonemal microtubules. The % of SPAG6+ cilia were reduced by 63.00 ± 4.082 (p-value = 0.0001, Student’s t-test) in the embryonic stage and by 73.67 ± 2.625 (p-value < 0.0001, Student’s t-test) in the postnatal stage. **(H)** *En face* images of embryonic respiratory cilia showing the central pair (CP) missing in *Kif27*^−/−^ mice taken using tissue ultrastructure expansion microscopy and the linescan across an axoneme. Scale bar = 2 µm. **(I)** Quantification of CP organization in the wt (n=340, N=3) and *Kif27*^−/−^ (n=340, N=3) mice for **(H)**. **(J)** Localizations for radial spoke (RSPH9 in cyan) and dynein arm (DNALI1 in cyan) proteins were unaffected in embryonic *Kif27*^−/−^ mice. Acetylated ⍺-tubulin in magenta labels the axonemal microtubules. Scale bar = 1 µm.

**Supplementary Figure 3.**
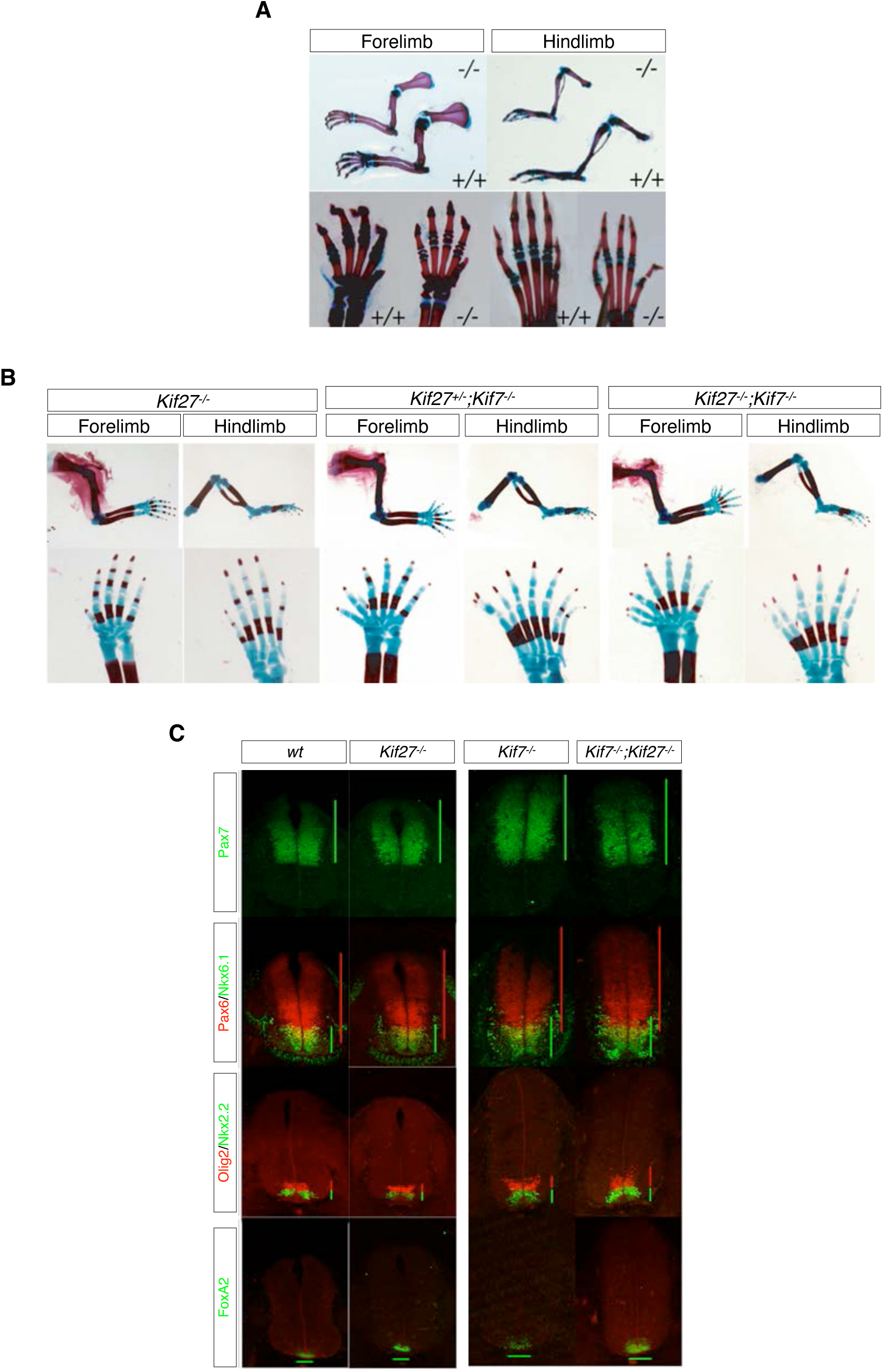
Hedgehog signaling is unaffected in *Kif27^−/−^* mice. **(A)** Alcian blue-alizarin red staining on P12 wt (‘+/+’) and *Kif27^−/−^* (‘−/−’) mice showed no apparent defects in skeletal patterning. Normal digit and limb patterning can be observed in both forelimb and hindlimb of the P12 *Kif27^−/−^* mutants when compared to the wildtype littermate. **(B)** Alcian blue-alizarin red staining on E18.5 wt and *Kif27^−/−^*; single mutants mice showed normal digit and limb patterning. E18.5 *Kif27^+/−^*; *Kif7^−/−^* and *Kif27^−/−^*; *Kif7^−/−^*mutants display severe polydactyly in both forelimb and hindlimb that are different from *Kif27^−/−^* single mutants and phenocopy the defects of *Kif7^−/−^* single mutants. **(C)** Neural tube patterning analysis was performed on *Kif27^−/−^*and *Kif7^−/−^* single mutants, as well as on and *Kif27^−/−^*; *Kif7^−/−^* double mutants. *Kif27* mutants exhibit comparable Pax7, Pax6, Nkx6.1, Olig2, Nkx2.2 and FoxA2 expression domain. Dorsal expansions for FoxA2, Nkx2.2, Olig2, Nkx6.1 were observed in both *Kif7^−/−^* single and *Kif27^−/−^*; *Kif7^−/−^* double mutants.

**Supplementary Figure 4.**
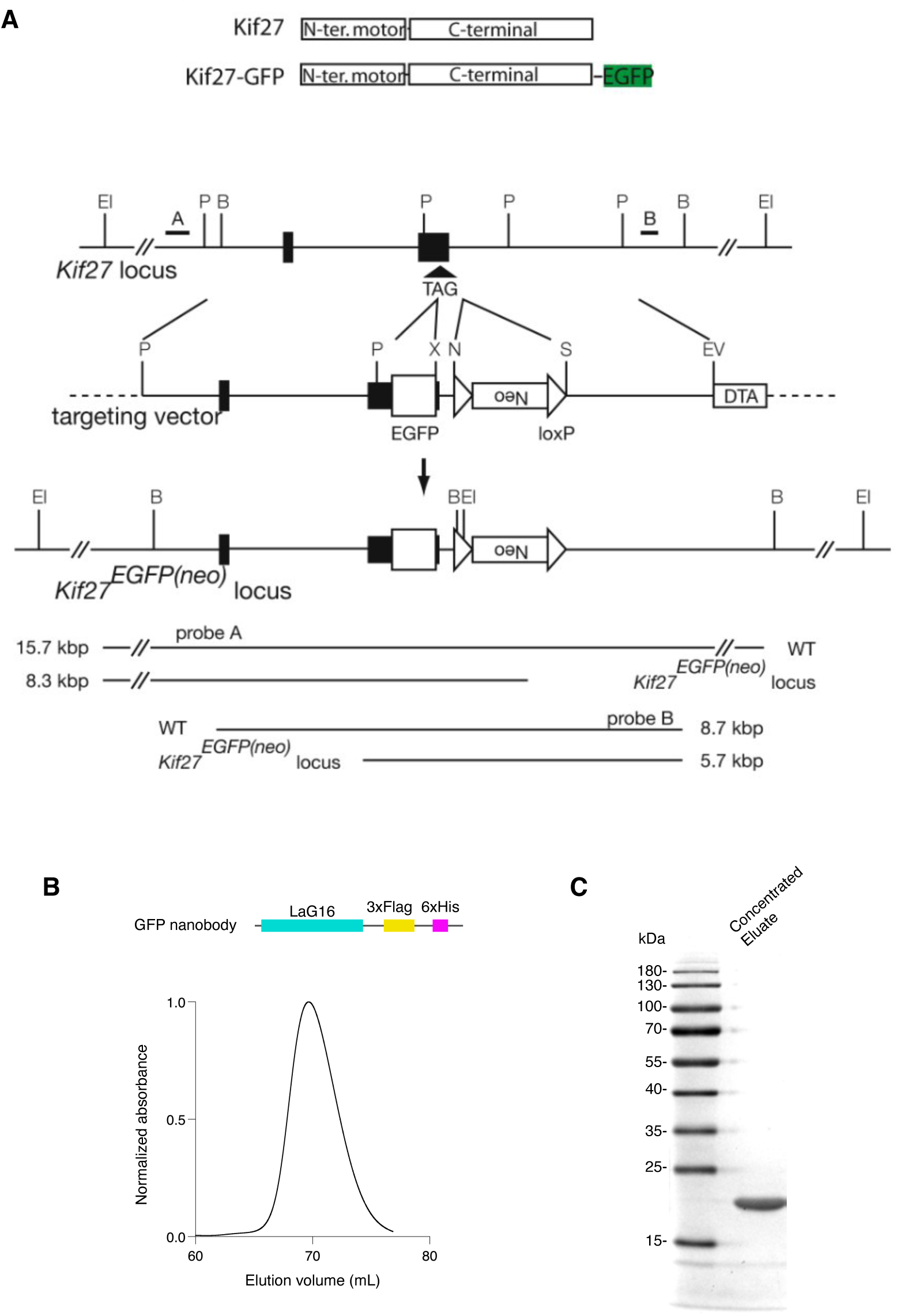
Generation of *Kif27^GFP/GFP^* mice and purification of anti-GFP nanobody. **(A)** Generation of the *Kif27^GFP^* alelle. The termination codon of *Kif27* was replaced with the EGFP coding sequence at the 3’ end of the last *Kif27* exon. The neo cassette was flanked with loxP sites (open arrowheads). Probes used for screening are indicated above the WT allele. The *Kif27^EGFP^*^(neo)^ allele was generated in ES cells by homologous recombination. Targeted mice were crossed with NLS-Cre mice, and the neo cassette was removed by Cre-mediated recombination. B; BamHI, EI; EcoRI, EV; EcoRV, N; NotI, S; SalI, X; XmaIII. **(B)** Design for the anti-GFP nanobody LaG16 coding region with 3xFlag and 6xHis tags at the C terminus and elution profile from size-exclusion chromatography of anti-GFP nanobody (peak volume, 69.66 mL). **(C)** Coomassie blue-stained SDS-PAGE analysis of purified GFP nanobody.

**Supplementary Figure 5.**
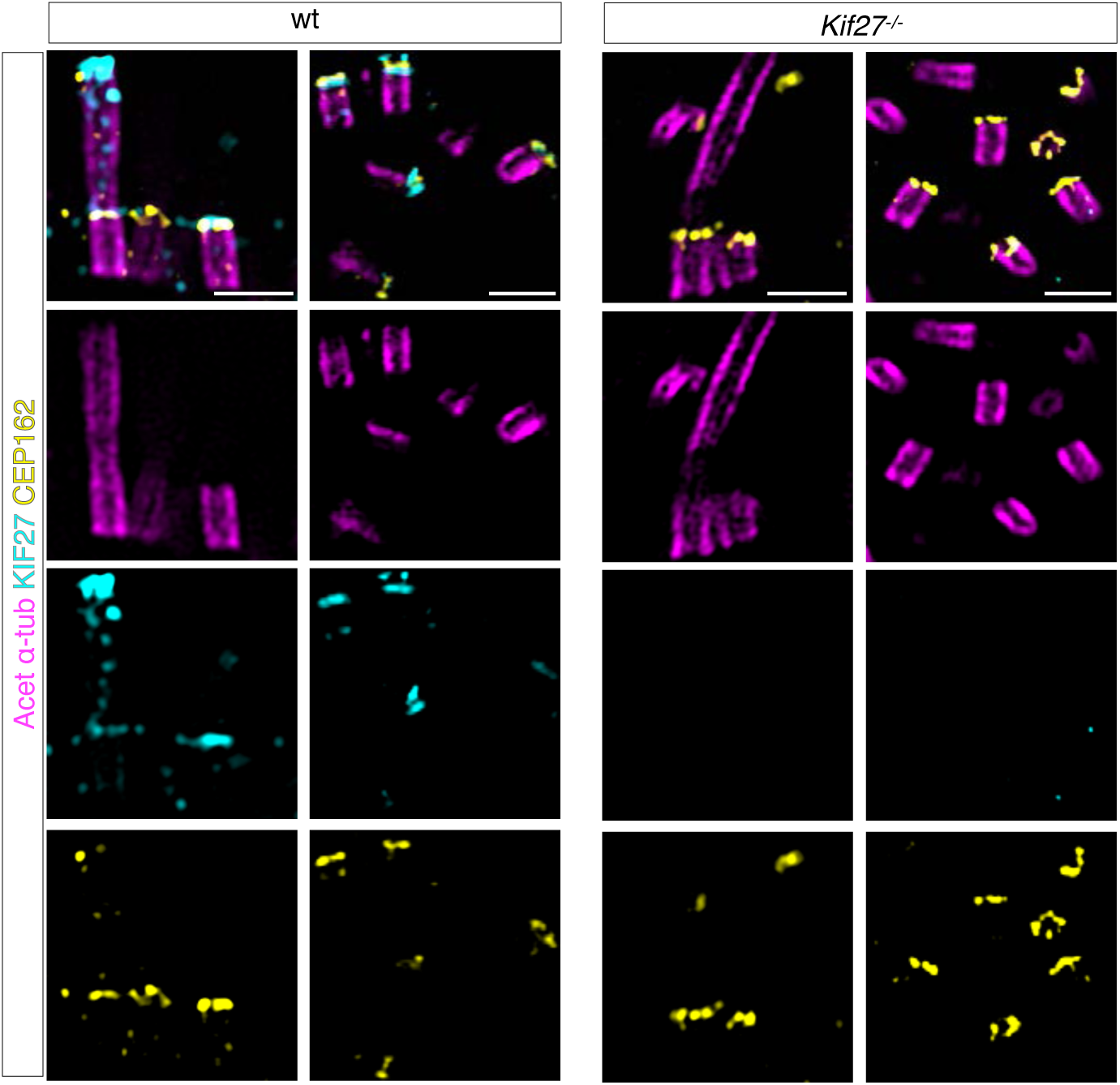
Validation of KIF27 antibody in expanded respiratory epithelia. Immunofluorescent staining using antibodies against KIF27 (cyan), CEP162 (yellow), and acetylated ⍺-tubulin (magenta) in expanded nasal epithelia of E17 wt and *Kif27^−/−^* embryos. Endogenous KIF27 co-localized with CEP162 at the distal centrioles, transition zone, as well as the tips of short cilia in wt embryos. KIF27 fluorescent signals were not detected in *Kif27^−/−^* embryos. Scale bars = 2 µm.

**Supplementary Figure 6.**
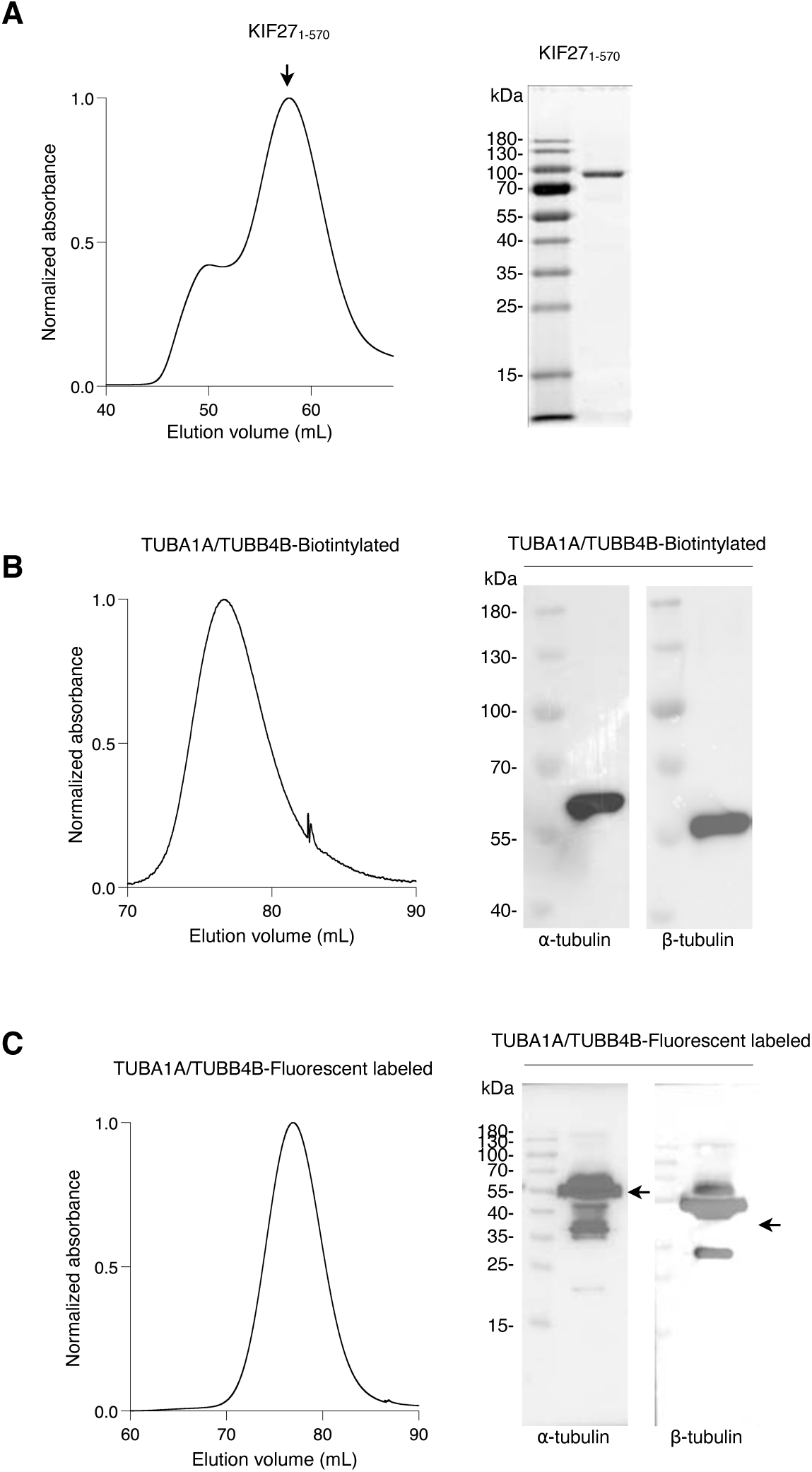
Purification of recombinant KIF27 proteins. **(A)** Elution profiles from size exclusion chromatography and Coomassie blue-stained SDS-PAGE of GFP-tagged KIF27_1-570_ (peak volume, 57.86 mL). Arrow indicates the expected position of purified protein. **(B and C)** Elution profiles from size exclusion chromatography and immunoblot analyses of biotinylated TUBA1A/TUBB4B (peak volume, 76.65 mL) **(B)** and X-rhodamine conjugated TUBA1A/TUBB4B (peak volume, 76.94 mL) **(C)**.

**Supplementary Figure 7.**
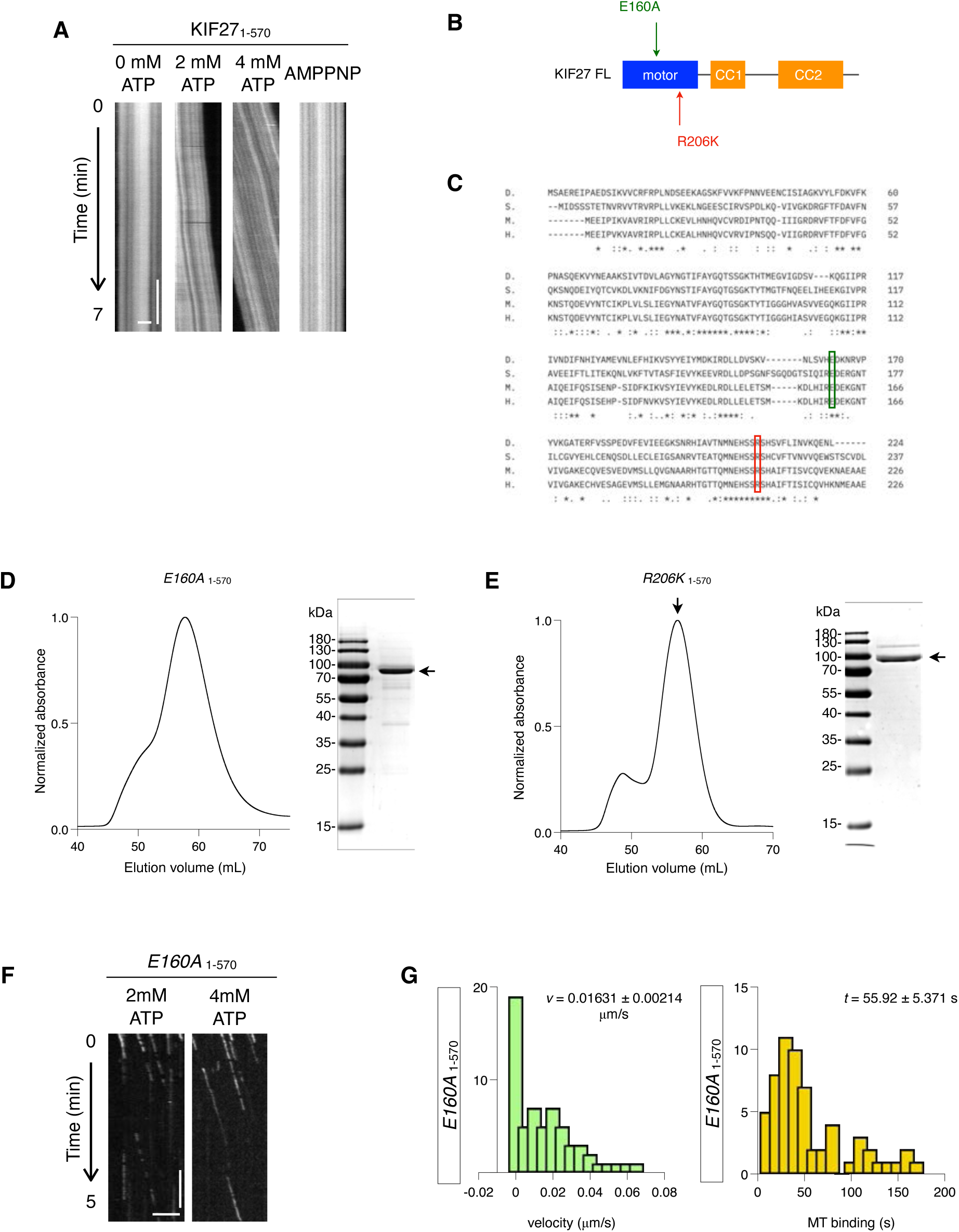
*In vitro* analysis of ATP-dependent KIF27 motility. **(A)** Representative kymographs from microtubule gliding assays showing polymerized microtubule (TUBA1A-TUBB4B) gliding driven by immobilized KIF27_1-570_-GFP. Horizontal scale bar = 2 µm. Vertical scale bar = 80 s. **(B)** Schematic of KIF27 domain organization and the locations of mutations tested. **(C)** ClustalW sequence alignment of *Drosophila* kinesin heavy chain (‘D.’, NCBI Reference Sequence: NP_476590.1), *Schmidtea mediterranea* KIF27 (‘S.’, SMED ID: SMED30001032, Planosphere), mouse KIF27 (‘M.’) and human KIF27 (‘H.’). Locations of conserved residues are indicated as boxes where point mutations are introduced,E160A in green and R206K in red. **(D and E)** Elution profiles from size exclusion chromatography and Coomassie blue-stained SDS-PAGE of KIF27^E160A^-GFP (peak volume, 56.50 mL) **(D)** and KIF27^R206K^-GFP (peak volume, 57.73 mL) **(E)**. **(F)** Representative kymographs from single molecule motility assays showing the single molecule motility of 20 pM KIF27^E160A^ on polymerized microtubule (TUBA1A-TUBB4B) in the presence of 2 mM or 4 mM ATP. Horizontal scale bar = 5 µm. Vertical scale bar = 60 s. **(G)** The average velocity and MT binding duration of KIF27^E160A^-GFP are shown as mean ± standard error of mean (n = 61).

**Supplementary Figure 8.**
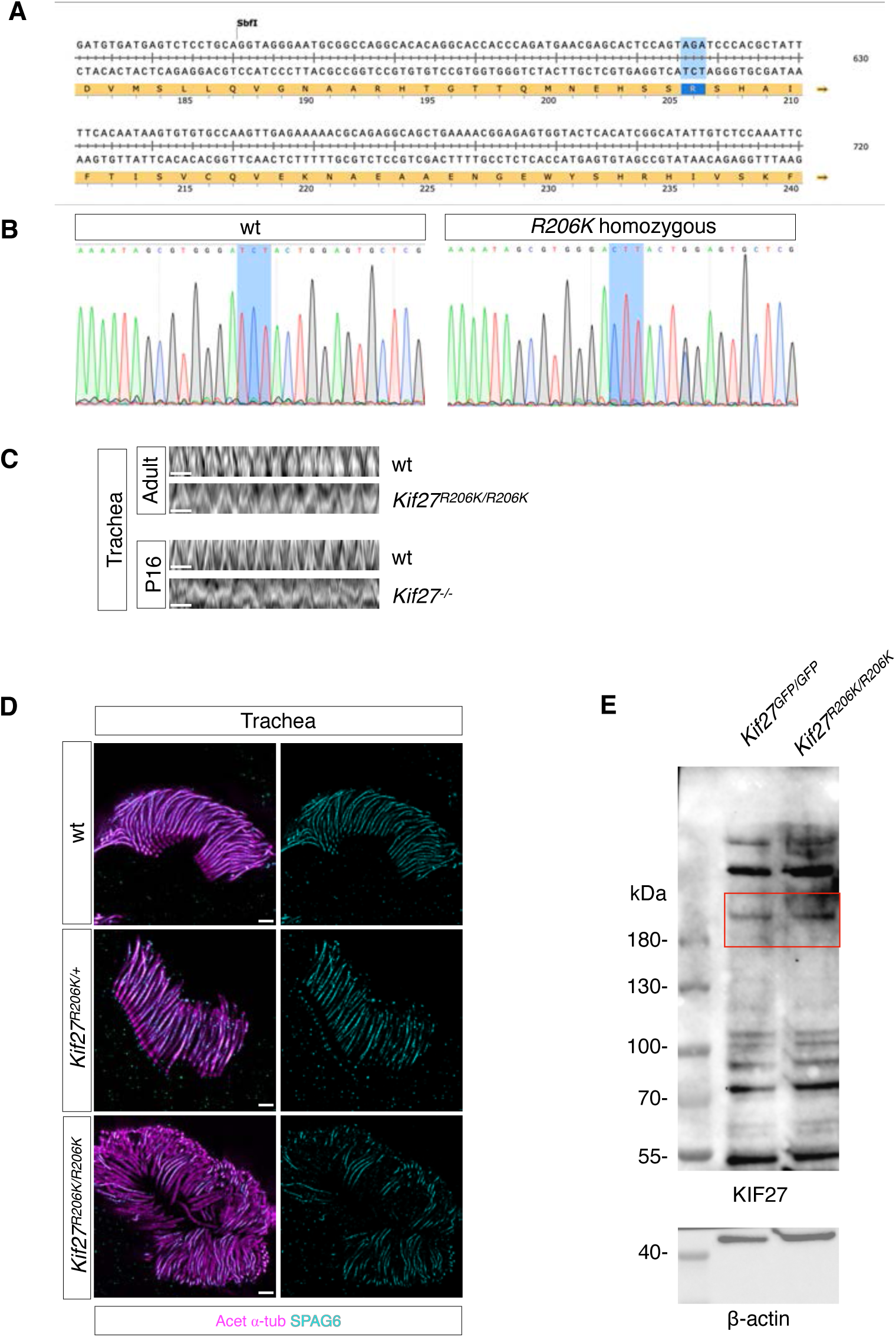
*Kif27^R206K/R206K^* mice show a rescued PCD-like phenotype compared to *Kif27^−/−^* mice. **(A)** DNA sequences for mouse *Kif27* spanning the targeted codon for R206 highted in red. R206 (AAG) was modified to R206K (AAG) **(B)** Chromatogram data for wt and *Kif27^R206K/R206K^* homozygous samples; a reverse primer 200bp downstream of the edited site was used for sanger sequencing. **(C)** Kymographs of beating motile cilia of adult wt and *Kif27^R206K/R206K^* mice trachea, and P16 wt and *Kif27^−/−^*mice trachea. Scale bar = 500 ms. Live imaging was captured at 30ms/frame for 30 seconds. **(D)** Sideview of tissue ultrastructure expansion microscopy images for wt, *Kif27^R206K/+^* and *Kif27^R206K/R206K^* mice trachea sections stained with acetylated ⍺-tubulin and SPAG6. Scale bar = 5 µm. **(E)** Immunoblots of anti-KIF27 antibodies using total cell lysates of *Kif27^GFP/GFP^*and *Kif27^R206K/R206K^* mTEC at air-liquid interface day 10. β-actin served as loading control. Regions of interest are highlighted by red rectangles.

**Supplementary Figure 9.**
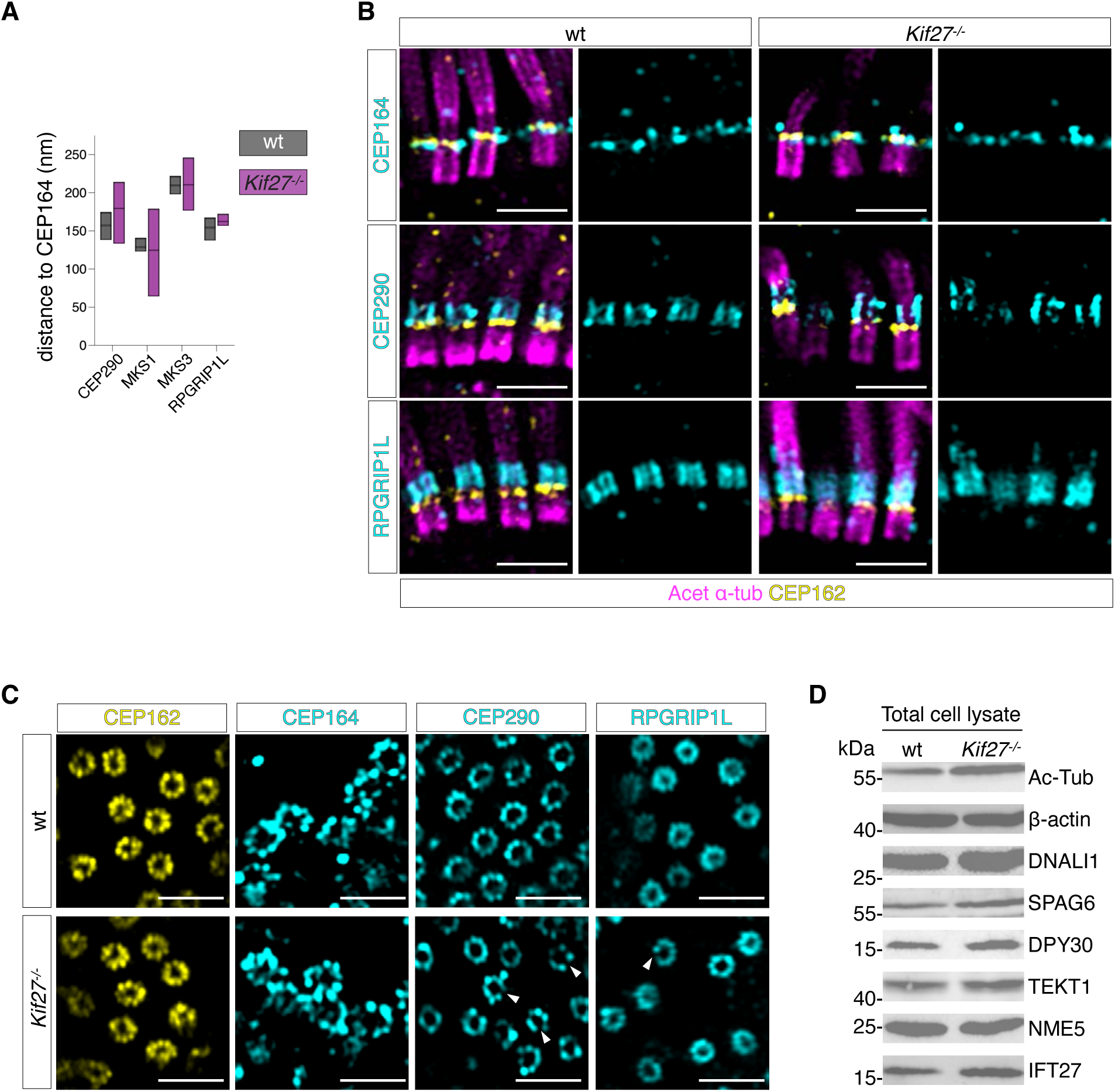
KIF27 is required for proper maintenance of the motile cilia transition zone. **(A)** Relative distance of different transition zone proteins from CEP164. For CEP290 distance for wt = 157.7 ± 18.39 nm and for *Kif27^−/−^* = 179.7 ± 41.70 nm. For MKS1 distance for wt = 129.2 ± 10.33 nm and for *Kif27^−/−^* = 124.9 ± 57.61 nm. For MKS3 distance for wt = 210.2 ± 12.05 nm and for *Kif27^−/−^*= 210.7 ± 34.77 nm. Lastly for RPGRP1L diastance for wt = 144.7 ± 12.30 nm and for *Kif27^−/−^*= 167.8 ± 8.198 nm. **(B)** Sideview images of tissue ultrastructure expansion microscopy images stained with CEP162 (yellow), Acetylated ⍺-tubulin (magenta), and CEP164, CEP290, and RPGRIP1L (all cyan) for wt and *Kif27^−/−^* mouse embryonic nasal respiratory cells. Scale bar = 3 μm. **(C)** *En face* images of tissue ultrastructure expansion microscopy images stained with CEP162 (yellow), CEP164 (cyan), CEP290 (cyan), and RPGRIP1L (cyan) for wt and *Kif27^−/−^* mouse embryonic nasal respiratory cells. Arrowheads indicate the gaps in the CEP290 and RPGRIP1L rings in *Kif27^−/−^* cilia. Scale bar = 3 μm. **(D)** Total cell lysates of wt and *Kif27^−/−^*mouse tracheal epithelial cells (mTEC) subjected to immunoblotting. Protein expression levels of ciliary proteins are unaffected in *Kif27^−/−^* cells. Acetylated α-tub marks cilia and β-actin served as loading control.

## Supplementary Movie S1-S4

Each movie is captured and displayed with 33.3 frames per second. Scale bar = 3 μm.

## Supplementary Table S1

*In silico* curation of potential regulators for motile cilia assembly. Gene symbols are shown as human homologues.

